# PRISM-G: an interpretable privacy scoring framework for assessing risk in synthetic human genome data

**DOI:** 10.1101/2025.10.17.682995

**Authors:** Alejandro Correa Rojo, Yves Moreau, Gökhan Ertaylan

## Abstract

Synthetic genomic data promises broader access, but unresolved privacy risks persist. In Europe, these risks increasingly hinder cross-border use of national genomic resources due to limitations in trust and legal interoperability rather than scientific demand. At the same time, privacy risk is not uniformly distributed: leakage driven by relatedness structure and rare-variant uniqueness can disproportionately affect underrepresented or vulnerable populations when shared data are misused or linked with external resources, making transparent, domain-aware measurement of privacy exposure central to responsible governance. We introduce PRISM-G, a model-agnostic framework that quantifies privacy exposure in synthetic genome data cross three complementary components: proximity to real genomes in genetic-coordinate space, replay of familial or population-structure patterns, and trait-linked exposure via rare variants and membership-inference signals. These components are normalized and combined through a risk-averse aggregation into a single 0-100 PRISM-G score. We evaluated PRISM-G on synthetic cohorts generated by a generative adversarial network (GAN), a restricted Boltzmann machine (RBM), and a logic-based SAT-solver (Genomator). Our results show that privacy vulnerabilities concentrate along different axes across models and marker densities, demonstrating that a single similarity-based metric is sufficient to characterize genomic privacy risk.

## Introduction

The increasing availability of large health data resources, such as biobanks and population cohorts, has accelerated biomedical discovery and clinical research by enabling advanced data analytics and the application of artificial intelligence at population scale ^1^. With declining DNA sequencing costs and the rise of population-scale genomics, genomic data has become central to studies of human variation and disease etiology, paving the way toward precision medicine and target therapeutics ^2,3^. However, the fact that personal genetic and health-related data is sensitive (Conduah et al., 2025; Evans & Jarvik, 2018; Shabani & Marelli, 2019), coupled to the fact that genomes are inherently re-identifiable ^7^, make broad data sharing a persistent technological, ethical, and legal challenge because demonstrated privacy risks and stringent regulatory requirements ^8,9^. This challenge is particularly acute in Europe, where the promise of cross-border genomic research and clinical translation is tightly coupled to governance, legal interoperability, and public trust. Multiple European initiatives, including the European Health Data Space (EHDS), aim to enable secure secondary use of health and genomic data at scale, yet operational data sharing has repeatedly stalled due to disagreements over acceptable risk, divergent national interpretations of data-protection obligations, and the lack of widely accepted technical evidence that released data cannot be linked back to individuals ^10–12^.

As a response to these challenges, synthetic data generation has emerged as a potential strategy to expand data availability while reducing privacy risks and preserving analytical utility without direct reliance on real personal records ^13^. In genetic research, artificial genome datasets are considered a promising tool, as many analyses depend on large-scale individual genetic variation that synthetic data could help provide for population genetics, rare disease studies, individual disease risk estimation, and improved representation of underrepresented populations ^14–16^. Moreover, synthetic data may facilitate genomic data sharing across collaborators and developers by easing regulatory constraints, while remaining subject to ongoing ethical and legal considerations ^17,18^.

A growing set of methods exists for generating synthetic genomic data. Traditional haplotype resampling combined with coalescent simulations (e.g., HAPNEST) provide scalable, reference-guided genotype synthesis and quality evaluation ^16^. More recently, deep generative models, such as generative adversarial networks (GANs), variational autoencoders (VAEs), restricted Boltzmann machines (RBMs), and diffusion models, aim to capture linkage disequilibrium, population structure and higher-order genomic dependencies ^19,20^. In parallel, logic-based generators (e.g., Genomator) formulate data synthesis as a constrained optimization problem to enforce biological plausibility and privacy constraints ^21^. Although these approaches show promise in producing faithful synthetic cohorts, generative modeling for genomics remains an active and maturing area of research, and synthetic genomic data is only beginning to emerge as a practical substrate for population genetics, method development, and downstream modeling (Battey et al., 2021; Wang et al., 2025).

Despite the demonstrated feasibility of synthetic genomic data generation, systematic evaluation of privacy risk remains a central challenge. Current practices rely mainly on similarity-based metrics, such as nearest-neighbor or Hamming-distance proximity between synthetic and real genomes, as proxies for identifiability ^16,21,24,25^, often complemented by Membership Inference Attacks (MIAs) that test whether individuals can be distinguished between training and holdout data ^26^. However, these approaches lack a unified interpretive framework and have sometimes led to the assumption that synthetic genomic data are “privacy-preserving by default”, despite evidence that disclosure risk varies substantially across methods ^27^. Moreover, models that accurately reproduce population-level distributions may still expose individuals to re-identification or membership detection through data linkage, genealogical inference, or trait prediction using external reference data and functional genomics ^8,28,29^. Consequently, the absence of standardized, domain-aware evaluation protocols limits meaningful comparison between generators and complicates risk-based data sharing ^30^, and regulatory assessment under legal frameworks such as the General Data Protection Regulation (GDPR) ^31^.

Genomic data are intrinsically multi-representational. The same individual can be described in coordinate space capturing population structure (e.g., eigenanalysis of common variants) ^32^, in kinship space reflecting familial relatedness through genetic relationship matrices (GRMs), and identical-by-descent (IBD) measures ^33^, and in trait-linked feature space summarizing biologically meaningful signals such as rare-variant burdens ^34^. Each of these representations encodes distinct aspects of personal information and reflects different ways in which individuals relate to populations, families, and traits.

### For this reason, privacy risk in synthetic genomic data cannot be assessed solely by how closely individual synthetic genomes resemble real ones

Even when individual-level similarity is low, privacy leakage may arise if synthetic cohorts preserve relational structure shaped by shared ancestry or reproduce feature-level signals linked to distinctive genetic characteristics. Evaluating privacy risk therefore requires accounting for multiple, interacting pathways through which personal information may be exposed, beyond simple distance-based measures.

Building on these representations we present the **Privacy Risk Integrated Score for Multi-representation Genomes (PRISM-G)**, a framework that evaluates synthetic cohorts across three complementary views and applies membership-inference tests based on rare variant content, calibrating the evidence into a single, interpretable risk score. PRISM-G summarizes privacy exposure via three representation indices: (1) **Proximity leakage (PLI)** asks whether any synthetic genome lies unusually close to a real (holdout) genome in genetic-coordinate space, closer than a normal population structure would explain; (2) **Kinship replay (KRI)** measures whether the synthetic cohort inadvertently recreates family structure or long-range dependence (e.g., close cousins, inflated relatedness, repeated haplotype segments); and (3) **Trait-linked leakage (TLI)** captures exposure that arises when simple membership inference signals or rare-variant burdens or collisions make individuals stand out. Each index yields a unit-interval risk (0–1), combined by a risk-averse “OR-like” aggregator and calibrated to a 0–100 scale using safe and leaky reference generators to ensure interpretability across datasets. A graphical description of the PRISM-G framework is illustrated in Figure 1.

**Figure 1.**
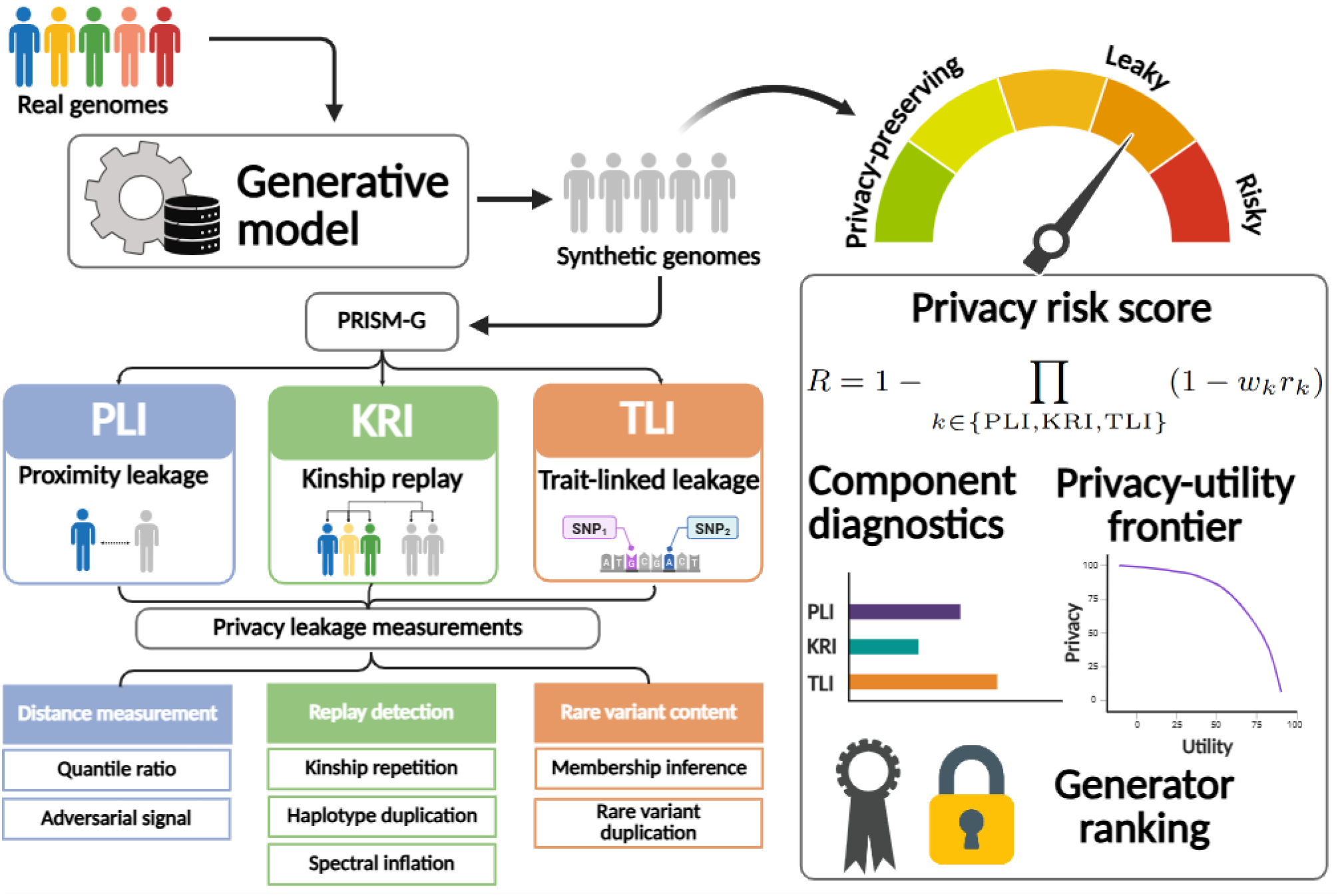
Overview of the PRISM-G privacy scoring framework.

**Figure 1.**
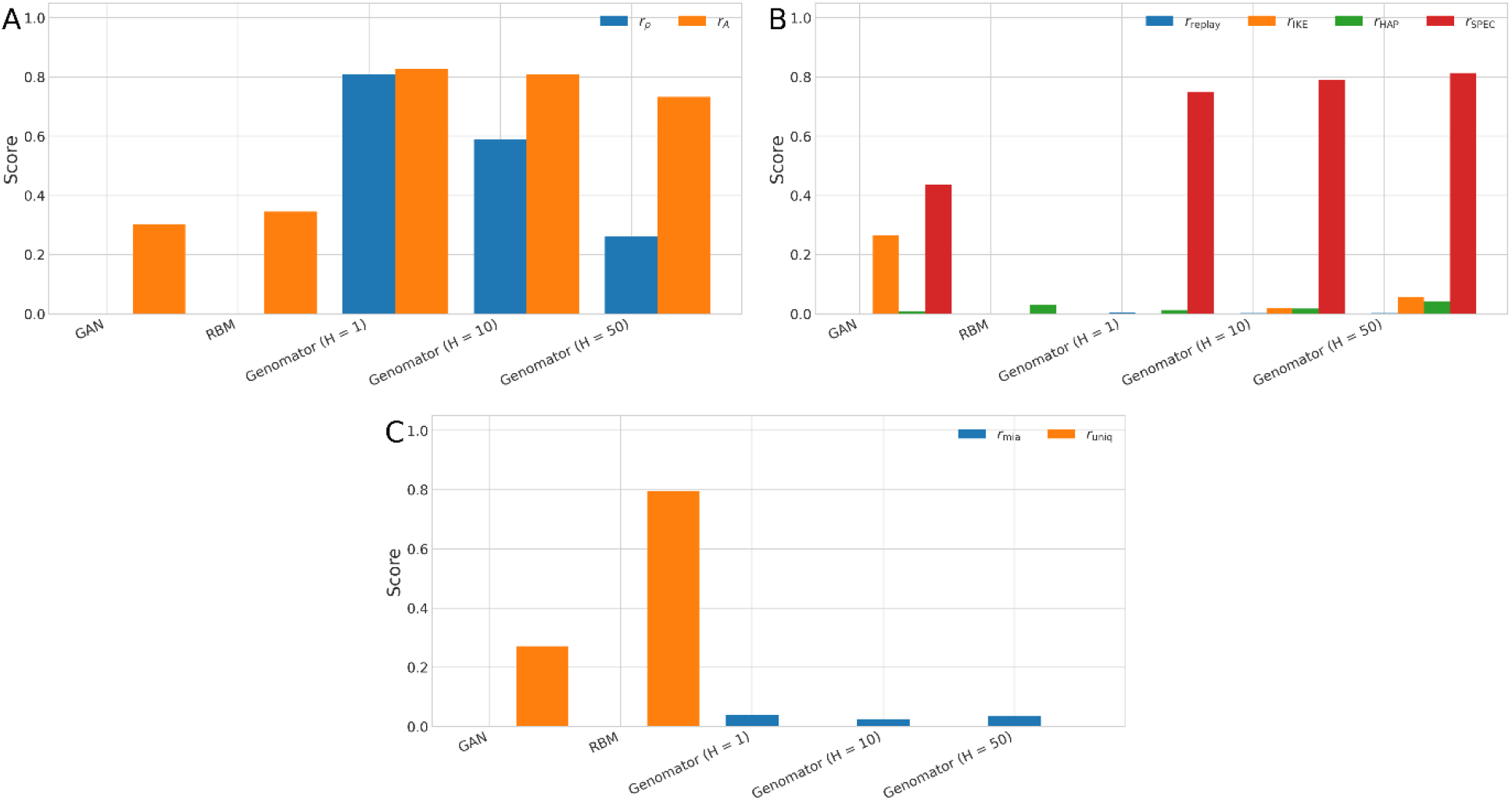
PRISM-G submetrics for the 10,000 SNPs panel dataset. Each barplot shows **A)** Proximity leakage components: proximity quantile and adversarial distance signals. **B)** Kinship replay components: replay, internal kinship excess, micro haplotype collision, and spectral anomaly. **C)** Trait leakage components: membership-inference performance, uniqueness and rarity match.

To demonstrate its applicability, we benchmarked three synthetic data generation approaches including GANs, RBMs, and methods based on SAT solvers, and evaluated their privacy risks across genomic representations, using subsets from the 1000 Genomes Project as reference dataset ^35^.

### Algorithmic basis and related work

As previously mentioned, early evaluations of synthetic genomic data often relied on similarity indices, such as distance to the nearest real neighbor, as proxy for identity risk. Yet genomic privacy risks are not limited to pointwise closeness: (1) population structure and relatedness create identifiable patterns that enable familial search and relationship inference, and (2) functional profiles and rare variants can reveal membership or traits even when overall distances look innocuous. Foundational work on principal component analysis (PCA) provides compact genetic axes that reveal population structure or demographic variation, while kinship estimation methods, such as GRM, operationalize relatedness at scale ^32,36,37^. In combination, these approaches can enable re-identification through distant relative matching in large databases. In parallel, privacy studies show MIAs against synthetic data and Beacon-style singling-out driven by rare variants ^27,38^. Taking this into account, PRISM-G’s components are designed to operationalize these documented risks using established genomic methodology: coordinates for proximity, GRM and short-haplotype patterns for relatedness replay, and MIA signals plus rare-variant counts for trait-linked exposure. Together, these components are aimed at capturing complementary and documented pathways through which personal information can leak from synthetic genome data. Table 1 shows an overview of the methodological basis and related work on the methods implemented in PRISM-G.

**Table 1.** Conceptual overview of the PRISM-G components, the genomic representations they probe, and the types of privacy leakage they are designed to detect. Formal definitions and implementation details are provided in the Materials and Methods section.

| Component | Algorithmic basis | PRISM-G adaptation | Adapted or based on |
| --- | --- | --- | --- |
| **Proximity leakage index (PLI)** | Projects genomes into PCA-derived genetic coordinates and tests whether holdouts are unusually close to synthetics in the lower tail of the distance distribution, benchmarked against holdout-to-holdout distances, with an adversarial nearest-neighbor check to avoid flagging ordinary structure. | Built on PCA for interpretable genetic axes, PRISM-G replaces raw nearest neighbor distance with a calibrated lower-tail closeness test and an adversarial sanity check, yielding a conservative proximity signal. | (Patterson et al., 2006; Yale et al., 2019.) |
| **Kinship replay index (KRI)** | Captures relatedness using GRM, tests for excess kinship against a safe baseline, scans for | PRISM-G combines four baseline-corrected signals: relatedness surplus, kin-edge | <sup>33,36,37,40–42</sup> |
|  | short haplotype sharing via short unphased genotype window, and flags spectral anomalies by detecting inflation of the GRM's leading eigenvalue. | replay, micro-haplotype collisions, and spectral concentration, into a novel aggregate that detects replayed familial structure and long-range dependence. |  |
| **Trait-linked leakage index (TLI)** | Build on rare-variant burdens to probe membership-inference signals in the synthetic cohort and estimates carrier probabilities from allele frequencies under Hardy-Weinberg equilibrium to flag excess co-occurrence of rare variants. | PRISM-G implements a simple nearest neighbor distance built from burden scores, and a rare-variant check that highlights unusual co-occurrence of uncommon variants. Together the results are mapped into a single score. | 26,34,38 |

### PRISM-G methodology

This section describes the formal definitions, estimation procedures, and calibration steps used to compute PRISM-G. We first define the three-component metrics, PLI, KRI, and TLI, each corresponding to a distinct genomic representation. We then describe how these components are aggregated and calibrated into a unified risk score, followed by robustness analyses for assessing sensitivity to calibration choices, and ranking risk for genomic data generators.

### Model description

Given a generative model, let *R*_tr_ denote the real training cohort of *m* individuals and *n* single nucleotide polymorphisms (SNPs) used to produce a synthetic cohort *S*, and let *R*_ho_ denote the real holdout cohort reserved for evaluation. PRISM-G evaluates *S* against *R*_ho_ in population coordinate space, kinship space, and trait linked feature space, respectively. We first describe the three components used to estimate PRISM-G. Each component produces a score in [0, 1], where 0 indicates no detected leakage on that axis and 1 indicates maximal exposure. Additional details on the methods used for component estimation can be found in the Supplementary Material.

### Proximity leakage index

The PLI metric asks whether any synthetic genome lies unusually close to a real individual in genetic coordinate space, beyond what would be expected from population structure alone. For computation, we project real and synthetic samples into a low-dimensional coordinate space using PCA of genetic variants. For each synthetic sample, we evaluate its distance to the nearest real individual and compare the resulting distribution to a reference baseline. Two complementary terms capture this signal: a lower-tail quantile ratio measuring over-closeness and an adversarial proximity score reflecting nearest-neighbor concentration. These terms are combined into a normalized PLI score as defined below.

Genotypes are embedded into a *d* dimensional space (principal components computed after standardizing genotypes using allele frequencies estimated from *R*_tr_) ^32^. Let 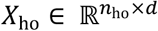 and 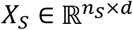 be the embedding for *R*_ho_ and *S*. For each holdout individual *i*, define the nearest synthetic distance as

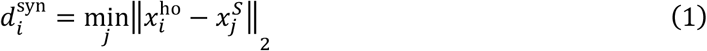

and the baseline nearest real distance to other holdout samples as

<H

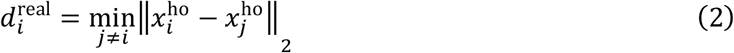

We compare the lower tails of the distance distributions estimated from Equations 1 and 2 via a quantile ratio:

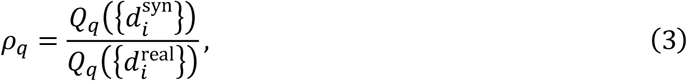

and define the proximity signal

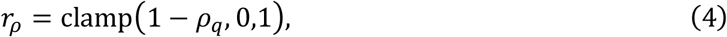

where clamp(*x*, 0,1) = min{1, max{0, *x*}}.

To add an adversarial check, we ask whether a holdout point is typically closer to another holdout than to any synthetic sample (Yale et al., 2019). Let 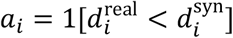 and 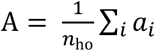. We map this to a risk as 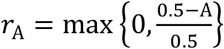. The PLI is calculated as

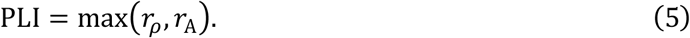

### Kinship replay index

The KRI metric evaluates whether synthetic datasets reproduce patterns of relatedness present in the real cohort, such as familiar relationships or population-level correlation structure. Unlike PLI, which focuses on individual proximity, KRI captures relational privacy risk that may persist even when no single synthetic genome closely matches a real individual. KRI aggregates several complementary diagnostics, each targeting a different manifestation of relatedness replay. These include replay of close kinship pairs, excess internal relatedness, haplotype-level collisions, and inflation of global correlation structure. We compute GRMs for *R*_ho_ and *S* after standardizing genotypes by training set allele frequencies as described by VanRaden, yielding *K*^ho^ and *K*^*S* 37^. KRI aggregates four signals, each baseline corrected against a safe or bootstrap reference.

First, the **replay** component (*r*_replay_) evaluates whether the synthetic dataset *K*^*S*^ reproduces the distribution of close-relatives relationships observed in the real cohort *K*^ho^. For a kinship threshold *θ*, we measure the similarity *M* between the real and synthetic close-kin spectra using the Jensen-Shannon (JS) divergence as measurement (Supplementary Material). We subtract a null rate *M*_0_ obtained by randomly permuting rows and columns of *K*^*S*^ and set

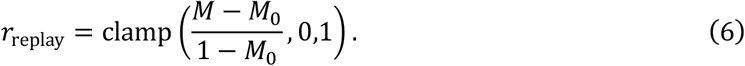

Second, the **internal kinship excess** (*r*_IKE_) tests for an overall surplus of relatedness in *S* (Supplementary Material). Let 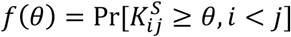. Using a holdout bootstrap set to estimate *f*_0_(*θ*), we define

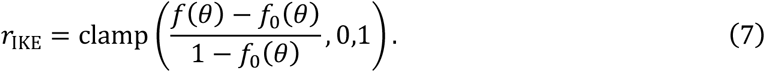

Third, the **micro haplotype collision** (*r*_HAP_) searches for excess reuse of short genotype patterns^42^. We slide fixed width windows of *k* SNPs and encode per window genotype patterns. For each pair of synthetic individuals, we count shared windows and compute a mean collision rate *c* (Supplementary Material). After subtracting a matched baseline *c*_0_, we set

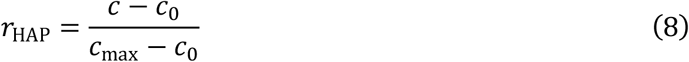

with *c*_max_ taken as the maximum rate across all windows.

Fourth, the **spectral inflation** (*r*_SPEC_) captures concentration of relatedness via the top eigenvalue of *K*^*S*^. Let *s* = *λ*_1_/trace(*K*^*S*^) and *s*_0_ be its baseline expectation (Supplementary Material), define

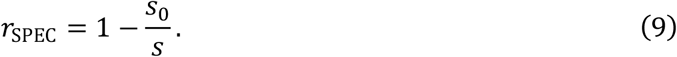

The KRI metric is estimated as

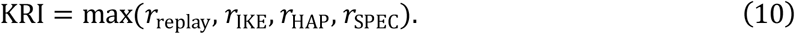

### Trait-linked leakage index

The TLI metric captures privacy risk arising from genetic features associated with phenotypes or distinctive individual characteristics, with a particular focus on rare genetic variants. Such features may enable attribute inference or membership attacks even in the absence of close genetic similarity. We quantify TLI using two complementary signals: performance of a simple MIA and the presence of rare-variant collisions between synthetic and real individuals. These terms reflect different ways in which trait-linked information may be exposed and are combined into a normalized TLI score as described below. Let *c*_*i*_ denote trait features for individual *i* (rare variant component burdens for chromosome sets)^34^.

For **membership inference advantage**, to probe whether training presence leaves a detectable trace in the synthetic cohort *S*, we construct a reference summary features from *c*_*i*_:

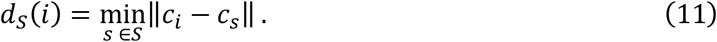

Using the *S*-referenced features, we score the training, and holdout sets with the negative distances *s*(*i*) = −*d*_*S*_(*i*) as membership scores, since being closer to *S* implies stronger membership evidence. We label the train individuals as 1 and holdout as 0, evaluate the area under the receiver operating characteristic (ROC) curve A and define

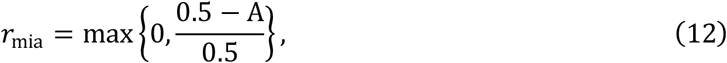

so random performance maps to zero and perfect separation maps to 1.

For **uniqueness and rarity match**, we evaluate whether rare signals occur more often in *S* than expected under a null hypothesis driven by training allele frequencies^38^. Define the rare set 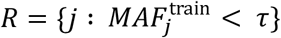. For SNP *j* with training allele frequency *p*_*j*_, under Hardy-Weinberg equilibrium approximation, the diploid carrier probability is *q*_*j*_ ≈ 2*p*_*j*_, and the null probability in a cohort of size *n*_*s*_ that at least two synthetic individuals carry the variant is

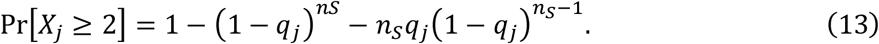

Let *U* be the observed fraction of rare sites with at least two carriers in *S* and *U*_0_ the average of the null probabilities above (Supplementary Material) and set

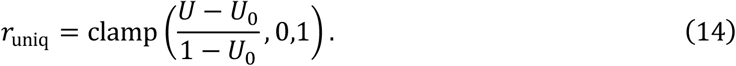

The TLI metric is estimated as:

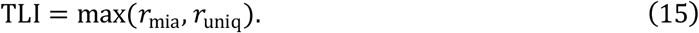

### Aggregation of component risks

Each component produces a unit-interval risk score with higher values indicating greater privacy risk: PLI for coordinate proximity, KRI for kinship replay, and TLI for trait linked exposure. Because raw component scores are not directly comparable across datasets or generators, we aggregate them into a single cohort-level risk score. To do so, we use a risk-averse, “OR-like” aggregation that preserves monotonicity and ensures that a single high-risk component cannot be masked by lower values in the others.

Specifically, let *r*_PLI_, *r*_KRI_, *r*_TLI_ ∈ [0, 1] denote the component scores and *w*_PLI_, *w*_KRI_, *w*_TLI_ ≥ 0 weights that sum to one. The aggregated raw risk is defined as:

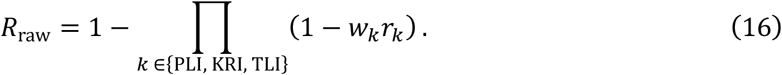

This construction has three practical advantages. First, it is **monotone** in every component: increasing any *r*_*k*_ increases *R*_raw_. Second, it is **risk-averse**: if any component approaches one, *R*_raw_ also approaches one regardless of the others. Third, it is **scale-free** with respect to cohort size, since components are already normalized to [0, 1].

When a sub-diagnostic inside KRI is not applicable (for example, the replay term when the holdout contains no close relatives), that term is omitted from KRI’s internal maximum without renormalizing other components. This preserves comparability across datasets.

### Calibration and estimation of weights

Because raw risk scores are not directly comparable across datasets or generators, PRISM-G uses simple reference generators to anchor the risk scale. This calibration step ensures that scores are interpretable and comparable without assuming a specific adversary model. We calibrate *R*_raw_ using two reference generators that anchor the lower and upper ends of the scale:

1. A **safe baseline** (*A*_Safe_) that preserves allele frequencies but intentionally removes dependence structure (for example, an allele-frequency–matched marginal sampler).
2. One or more **leaky baselines** (*A*_Leaky_) that intentionally overfit structure (for example, copycat, kinship-preserving variants).

To prevent the estimator from “spiking” on a single component and inflating overall risk, we calibrate the weights with a ridge-regularized fit to the anchor baselines. Concretely, we choose *w* to make the aggregated risk *R*(·; *w*) hit target values at the anchors while staying close to equal weights *w*_0_:

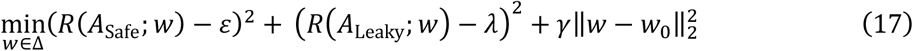

where the loss function is subject to *w*_*m*_ ≥ 0 and ∑_*m*_ *w*_*m*_ = 1.

Using Equation 17, we define three hyperparameters to calibrate the scale of *R*_raw_. The safe target *ε* anchors *R*(*A*_Safe_; *w*) to a small band (0.01–0.1), which avoids forcing any component weight to vanish purely to hit zero exactly. The leaky target *λ* (0.75-0.90) anchors *R*(*A*_Leaky_; *w*) high on the scale while avoiding saturation, leaving headroom for cases that are even leakier. Finally, the penalty strength *γ* (10^−4^ − 10^−2^) shrinks *w* toward the prior *w*_0_, balancing anchor fit with stability and discouraging spiky, single-component solutions.

### Estimation of PRISM-G score

For each baseline we compute the distribution of *R*_raw_. Let *α* denote the calibrated lower anchor of *R*_raw_ under the safe baseline, and let *β* denote the upper anchor, under the leaky baseline. We then map the observed raw risk for a model of interest to a standardized 0–100 scale:

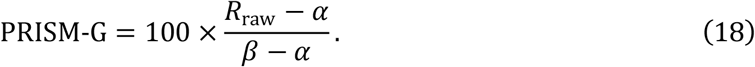

This linear calibration is monotone, model-agnostic, and yields scores that are comparable across datasets if the same anchors are used.

For interpretability, we group scores into qualitative bands: **Green, Amber**, and **Red**, indicating whether a method yields a safe, leaky, or risky dataset. These bands can be tuned to application-specific tolerances by shifting cutpoints or by defining them from anchor distributions.

Finally, to assess the robustness of the PRISM-G scores, we introduce a ranking stability test using Kendall’s rank correlation coefficient. Using bootstrapping, let *π*^(*b*)^ be the ranking at bootstrap draw *b*, and choose a baseline ranking *π*^(0)^. The stability test is defined as

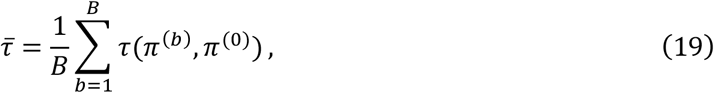

where 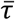 is the mean Kendall rank coefficient, and *B* the total number of bootstraps. The permutation value is obtained by comparing 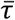 to *τ*(*π*^(perm)^, *π*^(0)^).

### Grid search evaluation

Finally, for hyperparameter tuning, we perform a grid search over the calibration parameters to maximize 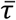 for PRISM-G score estimation. Given grids *ε* ∈ E, *λ* ∈ Λ, *γ* ∈ Γ, we evaluate each triple (*ε, λ*, Γ) and select the setting that yields the highest 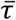:

1. Fit *w*^*^ by the objective function provided in Equation 17.
2. Compute PRISM-G for all candidate datasets/models.
3. Bootstrap anchors to obtain rank samples and compute Kendall’s 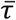 stability.
4. Return *ε, λ*, Γ that maximizes 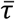.

### Estimation of the privacy-utility trade-off

The calibrated PRISM-G score provides a compact summary of privacy exposure, while component diagnostics reveal the mechanisms underlying potential leakage. Because generative models often trade fidelity and utility for privacy protection, PRISM-G can be paired with task-specific utility metrics to analyze privacy–utility trade-offs, for example, by generating a Pareto frontier. This enables systematic comparison between models and facilitates the identification of generators that achieve favorable balances between analytical utility and privacy protection.

### Evaluation procedure

#### Generative models

We evaluated PRISM-G on synthetic cohorts generated from three model families: (1) **GANs**, (2) **RBMs** as introduced by Yelmen and colleagues, and (3) **Genomator**, a logic-based generator that frames synthesis as a satisfiability (SAT) problem. Briefly, GANs pair a generator that maps random noise to genotype vectors with a discriminator that distinguishes real from synthetic samples: adversarial training drives the generator to match allele frequencies, population structure, and pairwise correlations, and conditional or convolutional layers provide control over ancestry and scalability. RBMs learn the joint distribution of genotype vectors via an energy function. After training (e.g., with contrastive divergence) they generate new genomes by alternating Gibbs updates, capturing linkage disequilibrium and higher order dependencies in the data^20^. In contrast, Genomator encodes admissible genotype configurations as logical constraints and uses SAT solving to sample synthetic genomes that adhere to specified biological and privacy rules, with a tunable parameter controlling distances to training data and within the synthetic cohort (Burgess et al., 2025).

As reference, we used the 1000 Genomes Project Phase 3 (1KGP) panel comprising 2,504 individuals. This served as the source for training/holdout partitions and for aligning markers across experiments.

#### Synthetic datasets

For the GAN and RBM baselines, we used the synthetic cohorts released by Yelmen et al. (2023), generated from 1KGP. For Genomator, we generated new synthetic cohorts from the same training splits under three Hamming distance settings *H* ∈ {1, 10, 50}, which regulate proximity between training and synthetic samples and among synthetic samples.

All analyses were performed on two datasets: one from Chromosome 15 with 10,000 SNPs and one from Chromosome 1 with 65,535 SNPs. For each dataset and generator, the synthetic cohort size matched the corresponding holdout cohort.

#### Parameter settings

For PRISM-G, each dataset was split into training (80%) and holdout (20%) sets. The training set was used to fit baselines for the component metrics and to define embeddings. The holdout set was used for final estimation of the component metrics and for membership-inference evaluation. The estimation of component metrics was performed over 1,000 bootstraps replicas.

For PLI metrics, we computed genetic coordinates using PCA on the training set and retained the first 10 components for all analyses. PLI used a fixed lower-tail quantile of 0.01 in the distance quantile ratio to focus on the closest real–synthetic pairs.

For KRI metrics, we used a kinship replay threshold of 0.125 (approximately first-cousin relatedness) with a tolerance *τ* = 0.01 when matching close-kin edges in the real data to those observed in the synthetic cohort. The *r*_IKE_ metric was evaluated at thresholds *θ ϵ* {0.10, 0.125, 0.25} and KRI took the maximum excess across these levels.

For TLI metrics, rare variants were defined by minor-allele frequency (MAF) less than 0.001, with allele frequencies estimated from the training set. Membership inference risk was computed as described in the PRISM-G methodology section.

For the initial estimation of raw values, we calibrated weights using hyperparameters *ε* = 0.02, *λ* = 0.75, and *γ* = 10^−3^, and performed ranking stability test over 1,000 bootstraps. Finally, for grid search, we evaluated *ε* ∈ {0.02, 0.04, 0.06, 0.08, 0.10}, *λ* ∈ {0.75, 0.80, 0.85, 0.90}, and *γ* ∈ {10^−4^, 10^−3^, 10^−2^} over 1,000 bootstraps.

#### Calibration anchors

We used two references to calibrate PRISM-G. The safe dataset is a sampler from a binomial distribution: for each SNP *j* with training-set allele frequency 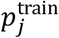, and for each synthetic individual *i*, we draw an independent diploid genotype:

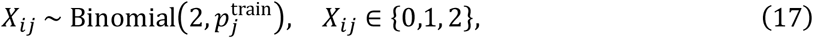

so that

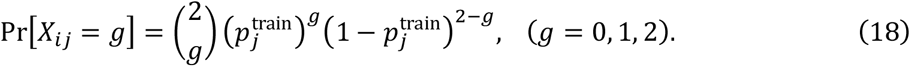

Draws are independent and identically distributed (i.i.d.) across *i* and *j*, using the same markers and sample size as the holdout set. This construction preserves site-level frequencies 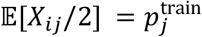 but removes linkage disequilibrium, kinship, and trait-linked correlations.

The leaky dataset is a copycat, kin-doped reference that intentionally overpreserves structure: each synthetic genome *X*_*ij*_ is templated to a training individual *t*(*i*) with small per-site perturbation (e.g., Hamming distance *H*(*X*_*i*_, *X*_*t*(*i*)_) ≤ *δ*), and a subset of templates is duplicated or paired to induce close-kin edges (target kinship *θ* in the range 0.25 – 0.125). This produces synthetic cohorts unusually close to training genomes and enriched for relatedness, serving as a high-risk anchor on the PRISM-G scale.

#### Utility evaluation and privacy-utility trade-offs

To evaluate the privacy-utility trade-off of the synthetic genomic datasets, we assessed their performance in a genetic ancestry inference task. Population structure was first estimated from the real training cohort using PCA. The first ten principal components were retained to define the genetic coordinate space used for downstream analyses. Synthetic genomes were then projected onto the same PCA embedding to ensure comparability between real and synthetic samples.

Utility was evaluated using the superpopulations labels provided by the 1KGP datasets, including African (AFR), European (EUR), East Asian (EAS), South Asian (SAS), and Admixed American (AMR) populations. A random forest classifier was trained on the real training samples using the ten principal components and the corresponding superpopulation labels and then applied to the synthetic datasets to predict ancestry assignments.

We quantified population-structure preservation using two complementary metrics. First, Procrustes similarity was computed between the centroids of real and synthetic ancestry clusters in the PCA space, providing a measure of geometric alignment between population structures. Second, we measured the JS similarity between the distributions of predicted ancestry labels in the synthetic data and the corresponding distributions in the real holdout data. JS similarity was calculated using the proportions of individuals assigned to each superpopulation group.

## Results

### Overview of PRISM-G components

We first evaluated submetric components for each synthetic dataset to illustrate how different generators exhibit distinct privacy leakage mechanisms prior to aggregation into PRISM-G scores. Figures 1 and 2, together with Supplementary Tables 1-4, summarize the results for the two marker panels. For the 10,000 SNP panel from Chromosome 15 (Fig. 1a; Supplementary Table 1), PLI submetrics indicated that GAN and RBM showed little proximity-based leakage, with quantile ratios above unity, yielding *r*_*ρ*_ = 0, and only modest adversarial proximity above chance (*r*_A_ < 0.35). In contrast, Genomator exhibited a tunable proximity profile (*H* ∈ {1, 10, 50}). Under tight constrains synthetic samples lay unusually close to holdout individuals, while increasing the constraint progressively reduced proximity-based exposure.

**Figure 2.**
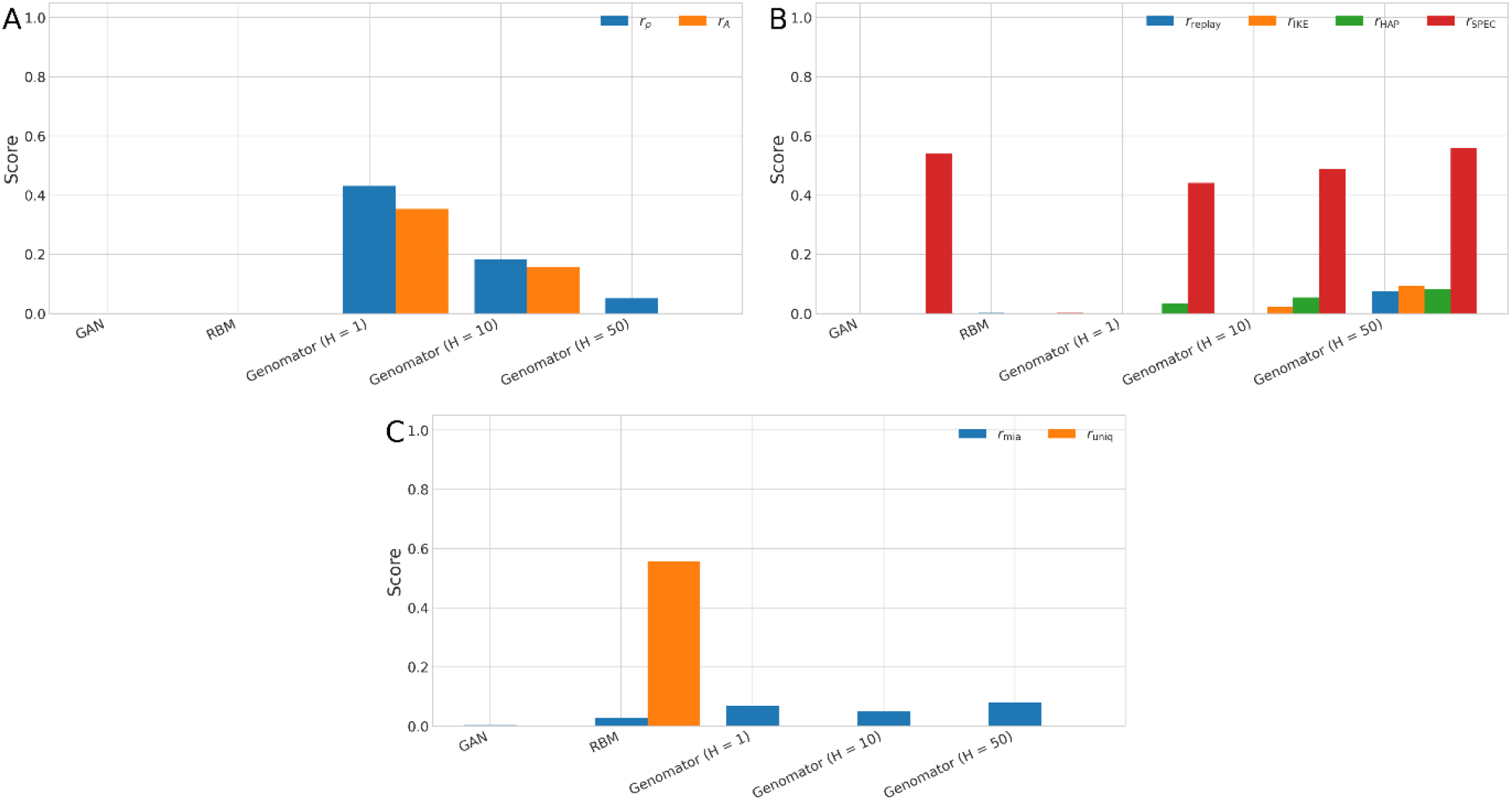
PRISM-G submetrics for the 65,535 SNPs panel dataset. Each barplot shows **A)** Proximity leakage components: proximity quantile and adversarial distance signals. **B)** Kinship replay components: replay, internal kinship excess, micro haplotype collision, and spectral anomaly. **C)** Trait leakage components: membership-inference performance, uniqueness and rarity match.

Kinship replay index submetrics (Fig. 1b; Supplementary Table 2) revealed distinct patterns across generators. Genomator’s kinship risk was driven primarily by spectral inflation (0.75 < *r*_SPEC_ < 0.81), with smaller contributions from the replay component (*r*_replay_), internal kinship excess (*r*_IKE_), and haplotype-collision terms (*r*_HAP_). GAN showed moderate inflation in *r*_SPEC_ and *r*_IKE_ whereas RBM exhibited minimal kinship-related signals across all submetrics.

For TLI (Fig. 1c; Supplementary Table 3), membership inference performance remained low for all models (*r*_mia_ < 0.04). However, rare variant uniqueness differed substantially: RBM showed the strongest rare-variant collision signal (*r*_uniq_ = 0.79), GAN exhibited moderate uniqueness (*r*_uniq_ = 0.27), and Genomator did not contribute across constraint setting (*r*_uniq_ = 0.00).

On the 65,535 SNP panel from Chromosome 1, PLI submetrics indicated little proximity-based leakage for GAN and RBM, whereas Genomator showed a monotonic reduction in proximity exposure as the constraint parameter *H* increased (Fig. 2a; Supplementary Table 4). In contrast, KRI submetrics revealed distinct generator-specific patterns. GAN exhibited moderate spectral inflation (*r*_SPEC_ = 0.54), RBM showed negligible kinship leakage across all KRI submetrics, and Genomator displayed a constraint-dependent shift, with haplotype-collision and spectral-relatedness signals (*r*_HAP_ and *r*_SPEC_) dominating at higher *H* values.

For TLI (Fig. 2c; Supplementary Table 6), GAN and Genomator showed low membership-inference signals (*r*_mia_) and negligible rare-variant uniqueness (*r*_uniq_) signals, whereas RBM exhibited elevated rare-variant uniqueness component (*r*_uniq_ = 0.56).

We next examined component-level PRISM-G scores to assess how proximity, relatedness, and trait-linked leakage contribute to overall privacy risk across generators. Figure 3 shows the radar plots of the component scores for each generator. For the 10,000 SNPs panel, GAN and RBM displayed moderate proximity leakage (0.30 < PLI < 0.35), with kinship replay moderate for GAN (KRI = 0.44) and low for RBM (KRI = 0.02). Trait-linked leakage was low for GAN (TLI = 0.27) but high for RBM (TLI = 0.79). Genomator presented high proximity leakage under tight constraints that decreased as constraints were relaxed (0.73 < PLI < 0.83), while kinship replay index remained elevated (0.75 < KRI < 0.81) and trait-linked leakage stayed low (0.02 < TLI < 0.04).

**Figure 3.**
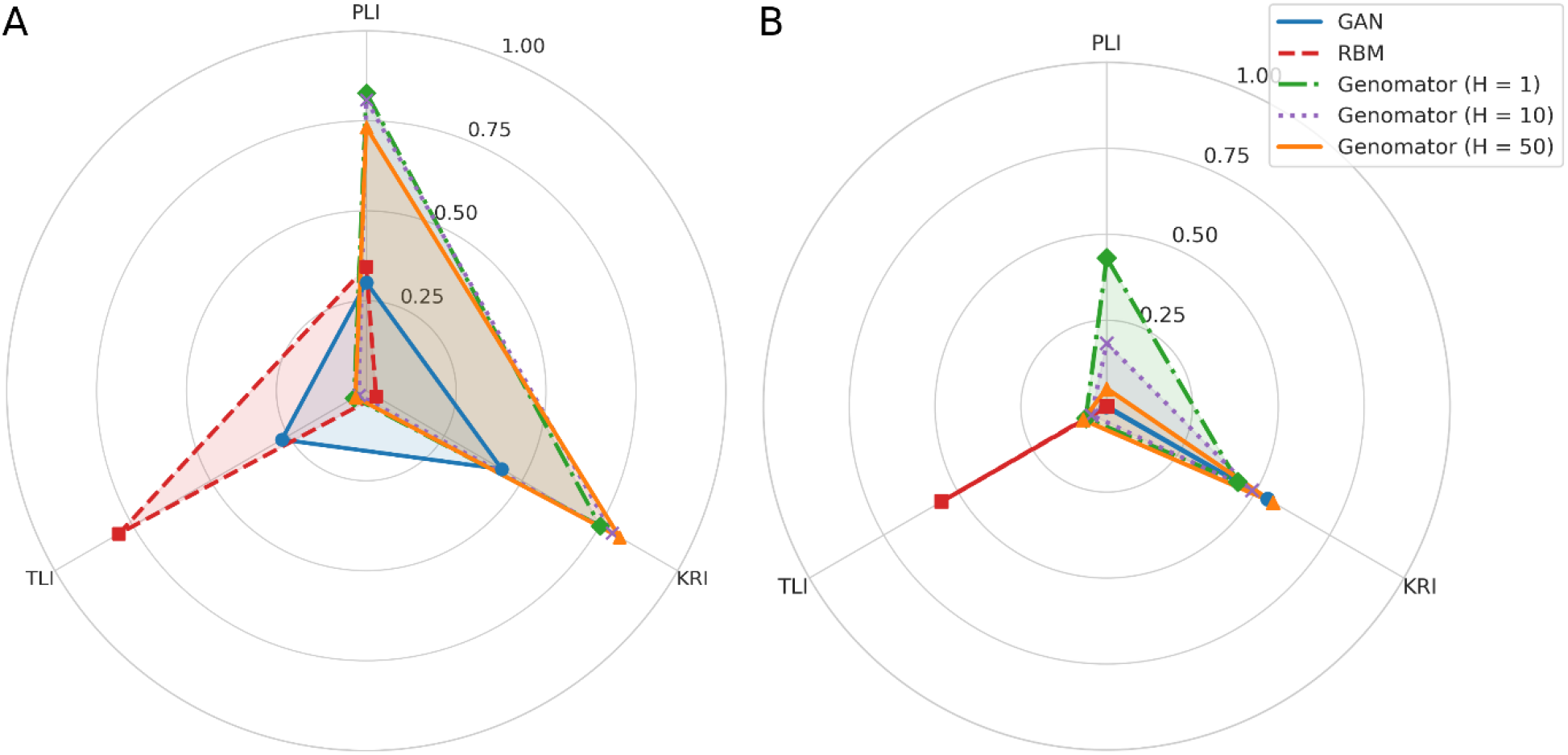
PRISM-G components. Each radar plot shows **A)** Metrics for the 10,000 SNPs panel dataset. **B)** Metrics for the 65,535 SNPs panel dataset.

On the 65,535 SNPs panel, GAN and RBM showed negligible proximity leakage. RBM displayed negligible kinship replay, whereas GAN showed moderate relatedness (KRI = 0.54). Trait-linked leakage remained minimal for GAN (TLI = 0.003) but elevated for RBM (TLI = 0.56). Genomator’s profile shifted toward lower proximity leakage with increasing constraint parameters (0.05 < PLI < 0.43), with intermediate kinship-related contributions (0.44 < KRI < 0.56) and low trait-linked leakage (0.05 < TLI < 0.08). Taken together, the component-level results show that privacy risk can emerge from different underlying mechanisms, including proximity over-closeness, relatedness inflation, and trait-linked exposure via rare genetic variants. Full component estimates are provided in Supplementary Tables 7-10.

### PRISM-G scores and ranking assessment

We next summarized the component-level privacy risk into a single PRISM-G score to enable compact comparison and ranking of synthetic genome data generators. To this end, we computed mean PRISM-G scores using bootstrapping (N = 1,000), calibrating each generator against safe and leaky reference datasets to place results on a common risk scale (see Methods). Figure 4 and Supplementary Tables 11-12 report the raw and calibrated PRISM-G scores for all generators on the two SNP panels.

**Figure 4.**
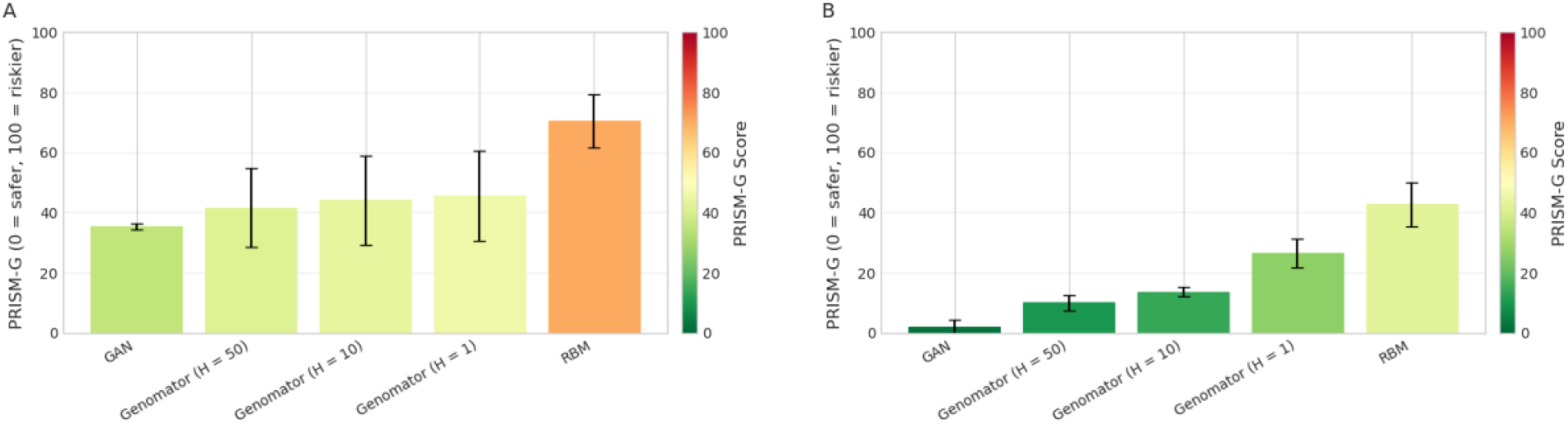
Mean PRISM-G scores for synthetic data generators with *ε* = 0.02, *λ* = 0.75, and *γ* = 10^−3^ as model hyperparameters. **A)** 10,000 SNPs panel datasets. **B)** 65,535 panel datasets.

On the 10,000 SNPs dataset from Chromosome 15 (Fig. 4a), the safe binomial sampler anchored the lower bound at 0 (*α* = 0.08) and the leaky copycat at 100 (*β* = 0.75). Among the learned models, GAN scored the lowest (PRISM-G = 35.31), while RBM scored the highest (PRISM-G = 70.37). Genomator yielded moderate scores on this panel, clustering between 41.58 and 45.53 across constraint settings, consistent with the elevated proximity and kinship leakage signals observed in the submetric components. Based on these scores, the generated datasets from GAN and Genomator were classified as **green** (safest; PRISM-G < 50), whereas the RBM-generated dataset was classified as **amber** (leaky; 50 < PRISM-G < 90) relative to the reference anchors.

On the 65,535 SNPs dataset from Chromosome 1, we observed the same relative ordering of generators (Fig. 4b). GAN again showed the lowest risk (PRISM-G = 2.10), while RBM yielded the highest score among the models (PRISM-G = 42. 76). Genomator’s scores decreased with looser constraints, spanning 10.11 to 26.53 across *H* settings. Using the same reference anchors as above, overall privacy risk scores on the denser panel were lower than those observed for Chromosome 15. In this setting, RBM remained the highest-risk generator, Genomator’s scores tracked its constraint *H* parameter, and GAN consistently showed low privacy risk. As a result, the datasets were labeled safe (PRISM-G < 50), at higher number of SNPs.

Using the bootstrap replicates, we then estimated Kendall’s rank correlation coefficient to assess how consistently the PRISM-G scores reproduced the model ranking within each dataset. Table 2 shows the summary statistics of the ranking stability test. Given the baseline ordering (RBM > Genomator > GAN; first bootstrap draw), the 65,535 SNPs dataset showed very high correlation (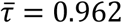 P-value < 0.018), whereas the 10,000 SNPs dataset showed moderate, non-significant correlation (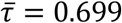 P-value < 0.074).

**Table 2.** Summary statistics of the Kendall ranking stability using *ε* = 0.02, *λ* = 0.75, and *γ* = 10^−3^ as model hyperparameters. SD: standard deviation, LCI: lower confidence interval, UCI: upper confidence interval.

| Dataset | $\bar{\tau}$ | SD | LCI (95%) | UCI (95%) | P-value |
| --- | --- | --- | --- | --- | --- |
| 10,000 SNPs |  |  |  |  |  |
| (Chromosome 15) | 0.699 | 0.287 | 0.200 | 1.000 | 0.074 |
| 65,535 SNPs |  |  |  |  |  |
| (Chromosome 1) | 0.962 | 0.081 | 0.800 | 1.000 | 0.018 |

### Sensitivity analysis and robustness evaluation

We evaluated the robustness of PRISM-G scores by performing a sensitivity analysis across a range of plausible calibration and regularization settings. In our previous results, PRISM-G was computed using a predefined set of hyperparameters *ε, λ*, and *γ*, which control anchor separation and regularization in the noisy-OR aggregation. We recomputed PRISM-G over a grid *ε* ∈ {0.02, 0.04, 0.06, 0.08, 0.10}, *λ* ∈ {0.75, 0.80, 0.85, 0.90}, and *γ* ∈ {10^−4^, 10^−3^, 10^−2^} at 1,000 bootstrap replicates per setting. Figure 5 shows the resulting mean aggregated scores. Across both SNP panels, PRISM-G decreased monotonically with increasing *λ*, while changes in *ε* primarily shifted score levels without altering relative ordering. The effect of ridge regularization varied with marker density, modulating the dispersion of scores without altering relative ordering. Overall, rankings and qualitative conclusions remained consistent across the grid, indicating that PRISM-G is robust to plausible variations in calibration and regularization parameters.

**Figure 5.**
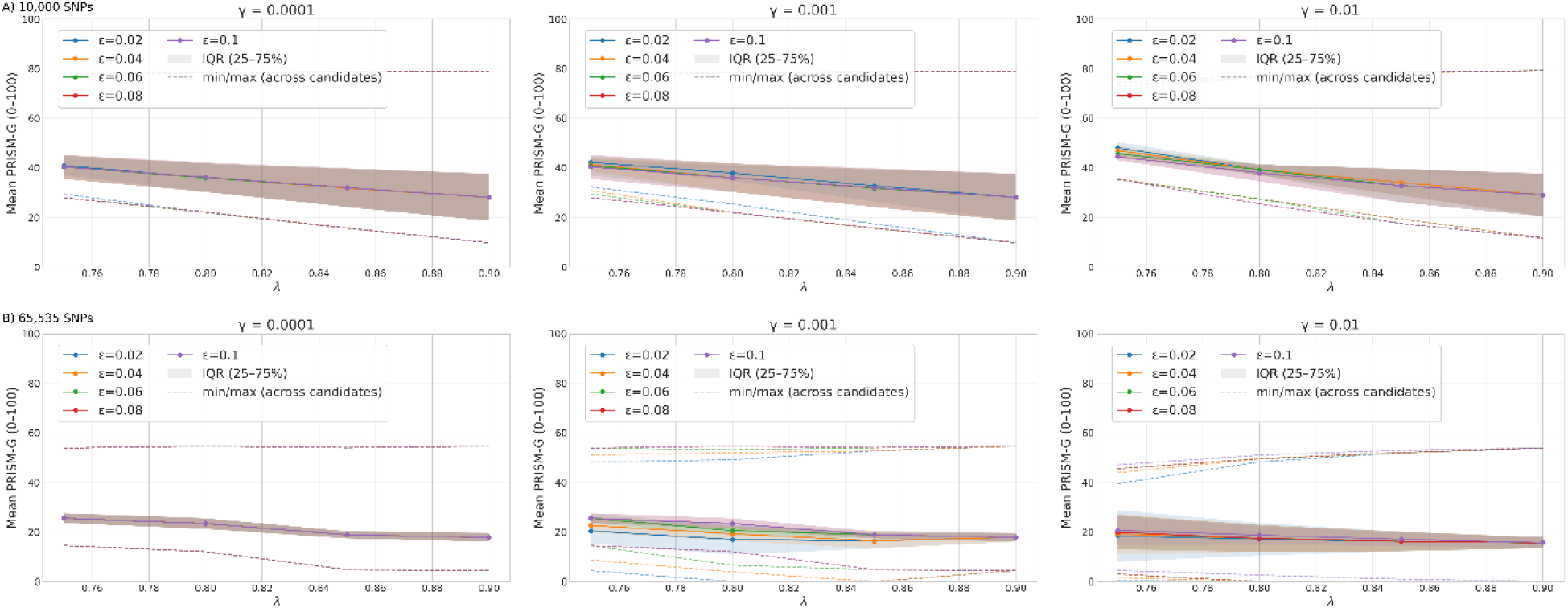
Sensitivity analysis plot to assess the estimated effect of model hyperparameters on PRISM-G scores. Mean PRISM-G scores are shown as a function of the calibration parameter *λ* for different values of *ε*. Shaded regions indicate the interquartile range of PRISM-G scores across bootstrap replicates, while dashed horizontal lines denote the minimum and maximum scores observed across replicates and hyperparameter settings. Panels show results for **a)** 10,000 SNPs panel datasets. **b)** 65,535 SNPs panel datasets. From left to right the plots correspond to *γ* ∈ {10^−4^, 10^−3^, 10^−2^}.

We performed a grid search evaluation to select the hyperparameters that maximizes 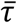. Table 3 and Supplementary Tables 13-16 shows the summary results and pairwise differences obtained for both datasets. With the best parameters obtained, the PRISM-G scores lead to a significant ranking with high Kendall’s correlation coefficients (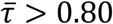; P-value < 0.018). Figure 6 shows the final estimated PRISM-G with significant pairwise comparisons across models. In both datasets, the same ranking was obtained despite differences in PRISM-G estimates.

**Table 3.** Summary statistics of the Kendall ranking stability using the optimal model hyperparameters obtained from grid search. SD: standard deviation, LCI: lower confidence interval, UCI: upper confidence interval.

| Dataset | $\varepsilon$ | $\lambda$ | $\gamma$ | $\bar{\tau}$ | SD | LCI<br>(95%) | UCI<br>(95%) | P-value |
| --- | --- | --- | --- | --- | --- | --- | --- | --- |
| 10,000 SNPs<br>(Chromosome 15) | 0.04 | 0.75 | 0.01 | 0.893 | 0.219 | 0.400 | 1.000 | 0.018 |
| 65,535 SNPs<br>(Chromosome 1) | 0.04 | 0.75 | 0.01 | 0.985 | 0.052 | 0.800 | 1.000 | 0.018 |

**Figure 6.**
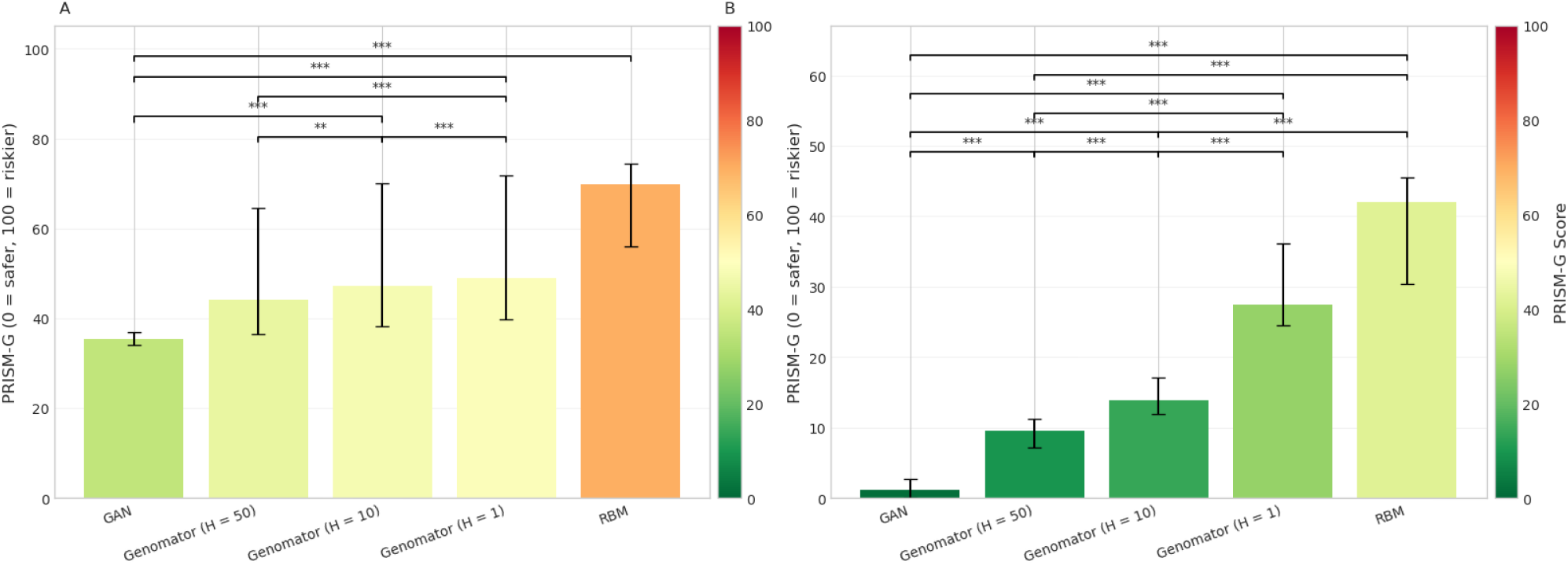
Mean PRISM-G scores for synthetic data generators and pairwise comparisons using the optimal model hyperparameters obtained using grid search. **A)** 10,000 SNPs panel datasets. **B)** 65,535 panel datasets.

### Privacy-utility analysis

Using the estimated PRISM-G scores as privacy metrics, we constructed a privacy-utility frontier to evaluate downstream analytical performance while controlling privacy risk. As a representative task, we assessed genetic ancestry prediction using population structure analysis as described in the Evaluation procedure section. Figure 7 shows the performance metrics obtained for both Chromosome datasets. Across generators, synthetic datasets preserved population structure and achieved high utility for ancestry prediction (Performance metric > 90%). Results on the PCA analysis for the synthetic datasets are shown in Supplementary Figures.

**Figure 7.**
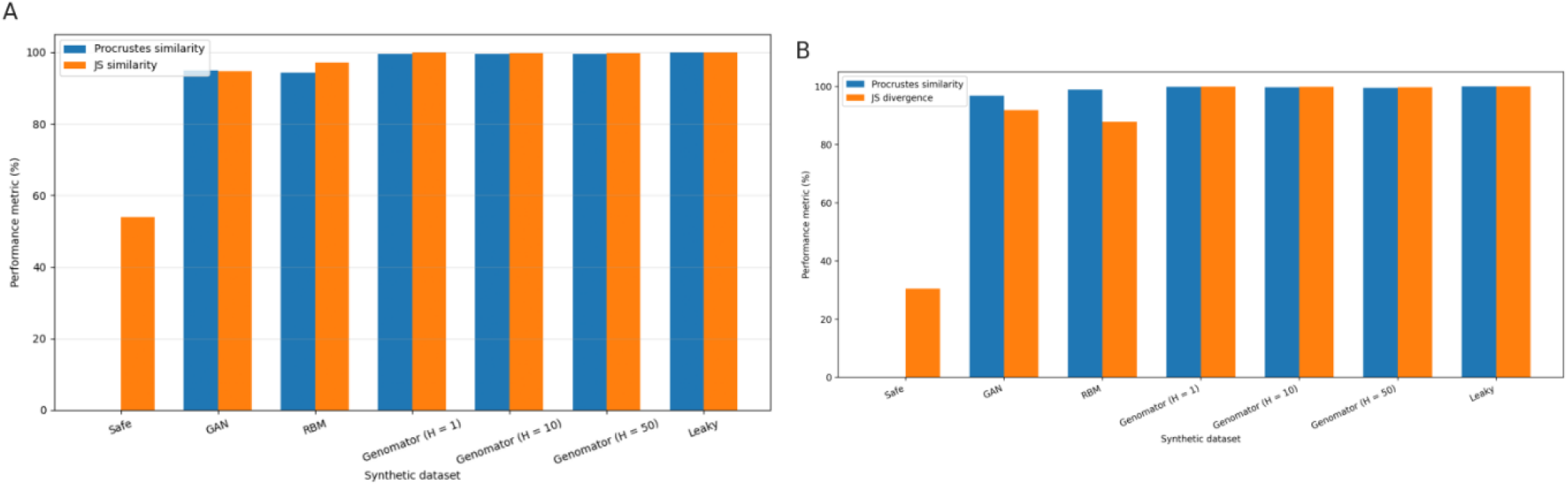
Performance metrics for population structure preservation and ancestry inference utility. **A)** 10,000 SNPs panel datasets. **B)** 65,535 panel datasets.

Pairing the PRISM-G scores with the average performance metrics as a utility measure for population structure analysis yields the privacy-utility frontier shown in Figure 8. The Pareto front represents the set of non-dominated solutions, where improving privacy would require sacrificing utility and vice versa. Genomator configurations lie on this frontier, maintaining high utility while reducing privacy risk as the Hamming constraint increases. GAN also achieves high utility with moderate privacy risks. In contrast, RBM is dominated, exhibiting higher privacy risk without a corresponding utility advantage.

**Figure 8.**
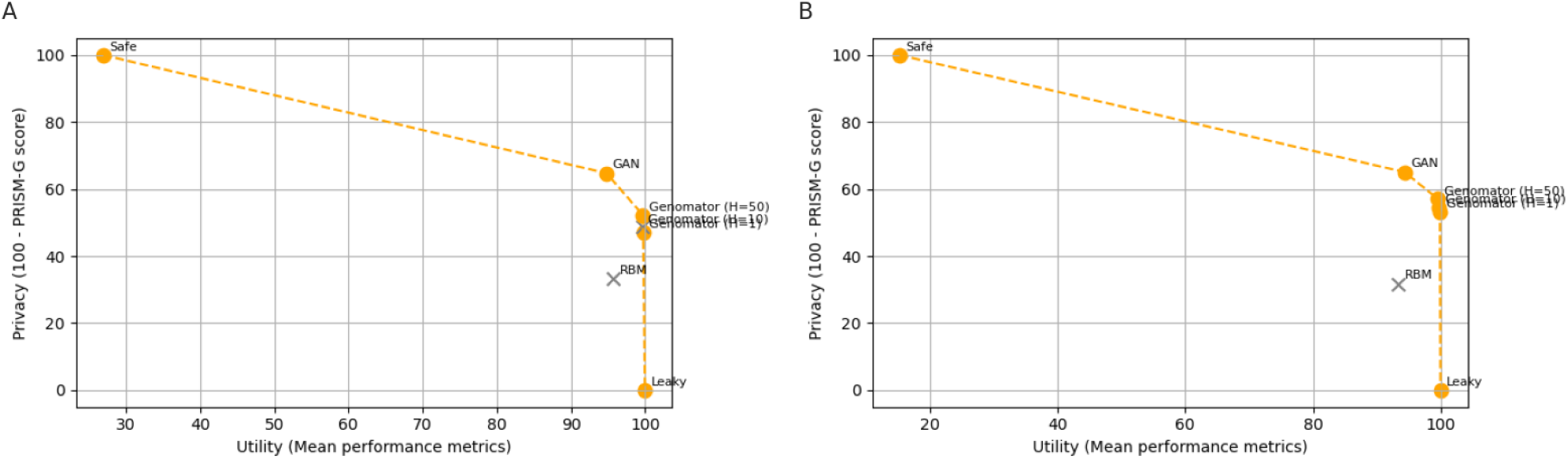
Privacy-utility trade-off curves. **A)** 10,000 SNPs panel datasets. **B)** 65,535 panel datasets.

## Discussion

In this study, we introduced PRISM-G, an integrated, model-agnostic framework for assessing privacy risk in synthetic genomic data across three complementary representations: genetic coordinate space (PLI), family structure (KRI), and trait-linked features derived from rare-variant content (TLI). Central to PRISM-G is the view that privacy risk in synthetic genomic data reflects potential leakage of personal information, which may arise not only from direct resemblance between synthetic and real genomes, but also from preserved relationships among individuals and from genetic signals linked to distinctive traits or characteristics.

Current evaluations of synthetic genomic data often lean on similarity-based diagnostics to argue that generated genomes are “far enough” from real sequences. However, such metrics have been shown to be insufficient as privacy guarantees, failing to address singling-out, linkability, and motivated-intruder scenarios ^30,44^. Moreover, genomic data encode personal information at multiple levels. Distance alone does not establish resistance to concrete attacks or capture risk arising from linked information ^45^: long-range familiar search in genealogy resources can re-identify individuals ^46^ and functional-genomics releases can leak attributes through cryptic channels and data linkage ^47,48^. PRISM-G makes distinct leakage pathways explicit, revealing failure modes that single-metric approaches may overlook. Our proximity component (PLI) offers a calibrated view of coordinate closeness. The kinship component (KRI) shows replayed relatedness and spectral concentration that can facilitate long-range familial search. The trait component (TLI) targets exposure through rare-variant collisions and simple membership-inference advantages. These components are recognized vectors for identification or singling-out ^38^. In doing so, PRISM-G aligns with and operationalizes emerging recommendations for domain-aware, task-linked evaluation and standardized reporting frameworks for synthetic medical data, moving beyond generic similarity indices ^49,50^.

Applying PRISM-G to multiple synthetic generators on two marker panels from the 1KGP dataset showed that privacy risk concentrates along different axes depending on model design and parametrization, as highlighted with prior attack-based evaluation of synthetic genomic data. Oprisanu and colleagues found that no single generator achieves both uniformly high utility and strong privacy and that some individuals remain vulnerable to membership inference even when population-level distributions are well matched ^27^. Using the GAN and RBM baselines from Yelmen et al. and a logic-based generator (Genomator) ^20,21^, PRISM-G component scores and submetrics reveal distinct patterns that reflect different modes of potential personal information leakage: at lower SNP density the GAN model exhibits a more balanced leakage profile, whereas at higher density it spreads samples sufficiently to avoid proximity and rare-variant collisions while still learning aspects of family structure. GANs are known to face mode-collapse that can produce synthetic data with limited diversity and, hence, hamper their effectiveness ^51^. However, methods and settings that improve coverage can mitigate collapse and increase sample diversity ^20,40^. Our observations align with improved coverage rather than collapse.

By contrast, our results suggest that RBM model tends to memorize rare variant patterns and family structures more strongly, amplifying trait-linked and familial leakage signals, thus elevating the corresponding PRISM-G components. Unintended memorization is a documented risk in generative models and can disproportionately affect uncommon samples, suggesting a plausible mechanism for our empirical findings ^52,53^. Aligned with our results, Genomator closely preserves global structure under tight distance constraints, elevating proximity and spectral-relatedness components while keeping rare variant content comparatively private. Relaxing the constraint reduces proximity leakage but can intensify spectral relatedness, so overall risk is panel-dependent, consistent with the developers’ observation that parameter choices modulate privacy risk.

Beyond interpreting PRISM-G and its components (PLI, KRI, TLI), Kendall’s rank stability test offers a robust way to compare models, indicating which one tends to produce safer, leaky, or risky datasets. Across both maker panels and all models evaluated, our results suggest that, for the 1KGP data, GAN produced the safest synthetic genomic dataset in PRISM-G terms, whereas RBM produced the riskiest. Taken together, our results reinforce that genomic “privacy” is inherently multi-faceted rather than a single-number attribute, with each model exhibiting distinct leakage patterns across genomic contexts that extend beyond simple distance metrics or rare-variant content.

Practically, three major key points emerge from our evaluation using PRISM-G. First, component scores are essential: the same integrated score can arise from different mechanisms, such as proximity over-closeness, inflated relatedness, or trait-linked uniqueness. Reporting PLI, KRI, and TLI metrics alongside the aggregate placement and statistical ranking can guide mitigation (e.g., tuning distance constraints to reduce PLI, decorrelating or enforcing pedigree pruning to reduce KRI, down-weighting rare-variant patterns to reduce TLI). Second, calibration against simple anchors yields an interpretable 0-100 scale without assuming a specific adversary model. In practice, we found these anchors bracketed the empirical range and made across-model comparisons straightforward. **Third, our results reinforce that no single generator is uniformly safest: risk shifts with marker density, cohort composition, and model hyperparameters**.

Apart from privacy exposure diagnostics, PRISM-G also enables joint evaluation of privacy and analytical performance through a privacy-utility frontier. Applying this framework shows that although all generators achieved similarly high utility for ancestry prediction their privacy risks differed substantially. Genomator configurations occupied favorable regions of the frontier, maintaining high utility while progressively reducing privacy risk as the Hamming distance constrained increased, whereas RBM exhibited higher privacy exposure without a corresponding gain in analytical performance. These results indicate that generators with comparable analytical utility may differ markedly in privacy risk.

Interpretable, component-level privacy auditing is essential for governance in multi-skateholder settings where trust depends on demonstrable safeguards. In European cross-border contexts, decisions about data sharing must be justified under heterogeneous legal interpretations and risk tolerances, making a single opaque similarity score insufficient ^10,12^. PRISM-G addresses this need by decomposing privacy exposure into proximity, kinship replay, and trait-linked (rare variant) mechanisms, enabling systematic auditing of synthetic genomic data and transparent comparison across generators. By making these mechanisms explicit, PRISM-G supports structured assessment of privacy risk and provides a basis for evaluating privacy-utility trade-offs for specific genomic tasks. In addition, the framework can support equity-aware evaluation, as proximity leakage related to ancestry structure and exposure driven by rare-variant uniqueness can vary across populations and may warrant additional scrutiny in future applications.

There are several limitations to note. PRISM-G does not provide a formal privacy guarantee (e.g., differential privacy). It is an empirical framework whose conclusions depend on chosen features, thresholds, and baselines. Our coordinate view used PCA on common variants. Alternative embeddings (e.g., UMAP or autoencoder latents) could change PLI sensitivity. KRI relies on GRM summaries and spectral patterns. Very long-range haplotype structure or phasing artifacts may modulate these signals. TLI focuses on rare-variant collisions and a minimalist MIA. Richer trait features (e.g., gene- or pathway-level burdens) or stronger attacks could alter rankings. Finally, our experiments were limited to subsets of 1KGP and two SNP panels. Applying PRISM-G to broader, more diverse population cohorts will better stress-test data generators and clarify how the metrics behave across ancestries, marker densities, and cohort designs.

Building on our findings, this work suggests several directions. Methodologically, refining the algorithms to yield robust scores for genome-wide synthetic panels, and extending TLI to task-specific phenotypes and functional genomics signals would better align privacy risk with real analytic use cases.

Empirically, large-scale tests on whole-genome data, with explicit admixture and rare-variant stratification, would stress the corner cases where replay and uniqueness are most likely. On the mitigation side, combining PRISM-G with privacy-preserving training or post-generation filters (e.g., data augmentation, allele-frequency-aware filtering, pedigree de-duplication, or differential privacy mechanisms) could translate component insights into concrete safeguards. Finally, frameworks such as PRISM-G could support broader auditing and governance workflows to provide structured evaluation of privacy risk, anonymization strategies, and data-sharing practices for future use of synthetic genomic data.

## Conclusions

Overall, PRISM-G offers a compact, calibrated ranking of privacy risk for synthetic genomic datasets while retaining interpretability across three complementary dimensions: PLI (proximity in genetic coordinate space), KRI (family structure and relatedness), and TLI (trait-linked features, particularly rare-variant exposure). Together, these components characterize where vulnerability arises, whether from over-closeness to real individuals, replayed kinship, or trait-driven uniqueness. Reporting both the integrated PRISM-G score and its component profiles is therefore informative: the single number supports deployment decisions and the choice of safeguards, while the submetrics explain why a model falls where it does on the 0–100 scale and point directly to targeted mitigations.

## Supporting information

Supplementary Results

## Author contributions

A.C.R. conceptualized the study, developed the methodology and conducted the experiments. G.E. and Y.M. supervised this work. The original draft was written by A.C.R. A.C.R., G.E., Y.M. reviewed and edited the draft.

## Declaration of generative AI and AI-assisted technologies in the writing process

The authors used ChatGPT to assist with reviewing the manuscript for grammar, clarity, and consistency, and to support debugging of code. Following the use of this tool, the reviewers and edited all generated outputs. The authors take full responsibility for the contents presented in this publication.

## Data availability

The 1000 Genomes Project is an open-access dataset available at https://ftp.1000genomes.ebi.ac.uk/vol1/ftp/release/20130502/. All synthetic genomes used in this study have been deposited in the PRISM-G repository on GitHub at https://github.com/alejocrojo09/prismg/tree/master/example_data.

## Code availability

Analysis scripts and codes are available on GitHub at https://github.com/alejocrojo09/prismg

