## Supplementary Material for "PRISM-G: an interpretable privacy scoring framework for assessing risk in synthetic human genome data"

**Supplementary Information**

**Replay metric**

The replay metric used in kinship replay index (KRI) evaluates whether the synthetic dataset replicates the distribution of close-relative relationships present in the real data. Let $K^{R}$ and $K^{S}$ denote the genetic relationship matrices computed for the holdout (real) data and the synthetic cohort $S$, respectively. From these matrices, we extract the upper-triangular kinship values:

$$\begin{aligned} R=\left\{ K_{ij}^{R}:i<j \right\}, S=\left\{ K_{ij}^{S}:i<j \right\}. \#\left( 1 \right) \end{aligned}$$

To focus on relatives rather than unrelated pairs, we restrict the analysis to the close-kin tail using a kinship threshold $\theta$:

$$\begin{aligned} R_{\text{close}}=\left\{ r\in R:r\geq\theta\right\},S_{\text{close}}=\left\{ s\in S:s\geq\theta\right\}. \#\left( 2 \right) \end{aligned}$$

In practice, $\theta=0.125$ corresponds approximately to second-degree relatives. The distributions of close-kin values are summarized using normalized histograms over the interval $[\theta, u]$ with $B$ bins. Let $P_{R_{\text{close}}}$ and $P_{S_{\text{close}}}$ denote the resulting distributions.

The replay metric is defined using the Jensen-Shannon divergence:

$$\begin{aligned} M=1-\frac{\text{JS}\left( P_{R_{\text{close}}},P_{S_{\text{close}}} \right)}{\log2}, \#\left( 3 \right) \end{aligned}$$

where $M \in[0,1]$ measures the similarity between the real and synthetic close-kin spectra, with higher values indicating stronger agreemeent.

To estimate the similarity expected under a null model, bootstrap samples are generated by resampling synthetic kinship values with replacement. Let $S^{(b)}$ denote the b-th bootstrap replicate and $M^{(b)}$ the corresponding replay similarity. The null expectation is defined as:

$$\begin{aligned} M_{0}=\mathbb{E}_{b}\left[ M^{\left( b \right)} \right]. \#\left( 4 \right) \end{aligned}$$

The final replay score is obtained by comparing the observed similarity $M$ to this null baselinie, yielding a measure of excess structural agreement between real and synthetic close-kin relationships.

**Internal kinship excess**

The internal kinship excess quantifies whether synthetic samples are mutually related than expected. Given $K_{\text{ho}}$ and $K_{\text{syn}}$, we define the fraction of pair exceeding a threshold $\theta$:

$$\begin{aligned} f\left( \theta\right)=\frac{\#\{\left( i,j \right):K_{\text{syn}}\left( i,j \right)\geq\theta, i<j\}}{\binom{n_{\text{syn}}}{2}} \#\left( 5 \right) \end{aligned}$$

To obtain the baseline expected fraction of related pairs, we estimate $f_{0}\left( \theta\right)$ by resampling from the kinship distribution of the holdout dataset. Specifically, we draw $m=\binom{n_{\text{syn}}}{2}$ kinship values at random, with replacement, from all pairwise kinship in the baseline matrix. For each bootstrap replicate $b$, we compute the proportion of sampled pairs whose kinship exceeds the thrseshold $\theta$:

$$\begin{aligned} f_{0}\left( \theta\right)=\frac{1}{m}\sum_{k=1}^{m} 1\left[ K_{0}^{(b)}\left( k \right)\geq\theta\right] \#\left( 6 \right) \end{aligned}$$

This process is repeated $B$, and the bootstrap average provides the expected baseline fraction:

$$\begin{aligned} f_{o}\left( \theta\right)=\frac{1}{B}\sum_{b=1}^{B} f_{0}^{\left( b \right)}\left( \theta\right) \#\left( 7 \right) \end{aligned}$$

The resulting internal kinship excess metric is then estimated as a normalized score above this internal null.

**Micro haplotype collision**

To quantify how often two or more synthetic genomes share the same short haplotype sequence, we compute the windowed haplotype collision across the synthetic genotype matrix $G_{\text{syn}}$. For a window of $k$ consecutive SNPs, we extract the genotype sub-matrix and encode each individual’s each individual genotype vector as a discrete key (haplotype string). The number of colliding pairs of identical haplotypes within that window is defined as:

$$\begin{aligned} C_{w}=\sum_{u} \frac{n_{u}\left( n_{u}-1 \right)}{2} \#\left( 8 \right) \end{aligned}$$

where $n_{u}$ is the count of samples sharing haplotype $u$. The per-window collision rate is then calculated as:

$$\begin{aligned} \rho_{w}=\frac{C_{w}}{\binom{n}{2}} \#\left( 9 \right) \end{aligned}$$

with $c_{\text{max}}= \max_{w} \rho_{w}$ as the maximum across all windows, and the overall collision rate for the dataset is obtained by averaging across all windows:

$$\begin{aligned} c= \frac{1}{W}\sum_{w} \rho_{w} \#\left( 10 \right) \end{aligned}$$

To derive baseline expectation, the same statistic is computed on the holdout genotype matrix over bootstrap replicates:

$$\begin{aligned} c_{0}=\frac{1}{B}\sum_{b=1}^{B} c_{0}^{\left( b \right)} \#\left( 11 \right) \end{aligned}$$

**Spectral inflation**

The spectral inflation component detects abnormal concentration of relatedness or latent clustering in the synthetic kinship matrix. Given $K_{\text{syn}}$, we approximate its largest eigenvalue $\lambda_{1}$ using power iteration:

$$\begin{aligned} v_{t+1}=\frac{Kv_{t}}{\left\| Kv_{t} \right\|}, \lambda_{1}=\lim_{t\to\infty} \frac{\left( Kv_{t} \right)\cdot v_{t}}{v_{t}\cdot v_{t}} \#\left( 12 \right) \end{aligned}$$

We then compute the normalized leading-eigenvalue share:

$$\begin{aligned} Type equation here.s=\frac{\lambda_{1}}{\text{trace(}K\text{)}} \#\left( 13 \right) \end{aligned}$$

where $\text{trace(}K\text{) = }\sum_{i} K_{ii}$ is the total diagonal sum of the kinship matrix, representing the total genetic variance across all individuals. A large value of $s$ indicates that a single eigenmode captures a disproportionate share of total variance, reflecting over-clustered or replayed structure in the synthetic cohort.

To derive baseline expectation, the same statistic is computed on the holdout kinship matrix $K_{\text{ho}}$:

$$\begin{aligned} s_{0}=\frac{\lambda_{1}\left( K_{\text{ho}} \right)}{\text{trace(}K_{\text{ho}}\text{)}} \#\left( 14 \right) \end{aligned}$$

**Uniqueness and rarity match**

The uniqueness and rarity match used in trait-linked leakage index (TLI) quantifies whether rare variants appear more frequently in the synthetic data than expected by chance. For each SNP $j$with allele frequency $p_{j}Type equation here.$, we approximate the diploid carrier probability under Hardy-Weinberg equilibrium as $q_{j}\approx2p_{j}$. Given a synthetic cohort of size $n_{s}$, let $X_{j} \sim\text{Binomial}(n_{s},q_{j})$ denote the number of synthetic individuals who carry at least one copy of the variant.

To compute the probability that at least two synthetic individuals carry the variant by random chance, we use:

$$\begin{aligned} \Pr\left[ X_{j}\geq2 \right]=1-\Pr\left[ X_{j}=0 \right]-\Pr[X_{j}=1]\#\left( 15 \right) \end{aligned}$$

Under the binomial model, we then define:

$$\begin{aligned} \Pr\left[ X_{j}\geq2 \right]=1-\left( 1-q_{j} \right)^{n_{s}}-n_{s}\left( 1-q_{j} \right)^{n_{s}-1} \#\left( 16 \right) \end{aligned}$$

This term represents the null expectation that a rare allele seen in the training data could appear at least twice in a random sample size $n_{s}$, assuming independent draws from the allele-frequency distribution. Averaging this probability across all rare variants yields $U_{0}$, the expected fraction of rare sites with $\geq$ 2 carriers under the null. From the synthetic dataset we compute $U$, the observed fraction of rare variants that have at least two carriers:

$$\begin{aligned} U=\frac{1}{\left| R \right|}\sum_{j} 1\left[ X_{j}\geq2 \right] \#\left( 17 \right) \end{aligned}$$

If the synthetic data inadvertently replays or memorizes rare variants from the training cohort, $U$ will exceed $U_{0}$.

$$Type equation here.$$
