## Supplementary Results for "PRISM-G: an interpretable privacy scoring framework for assessing risk in synthetic human genome data"

**Supplementary Tables**

| **Model** | **ρ_q_** | **r_ρ_** | **A_adv_** | **r_A_** |
| --- | --- | --- | --- | --- |
| GAN | 1.053 | 0.000 | 0.349 | 0.301 |
| RBM | 1.099 | 0.000 | 0.327 | 0.345 |
| Genomator (H = 1) | 0.191 | 0.809 | 0.086 | 0.828 |
| Genomator (H = 10) | 0.412 | 0.588 | 0.096 | 0.808 |
| Genomator (H = 50) | 0.738 | 0.262 | 0.134 | 0.733 |

**Supplementary Table 1.** Proximity leakage index (PLI) submetrics for the 10,000 SNPs panel dataset (chromosome 15).

| **Model** | **r_replay_** | **r_IKE_** | **r_HAP_** | **r_SPEC_** |
| --- | --- | --- | --- | --- |
| GAN | 0.000 | 0.266 | 0.008 | 0.435 |
| RBM | 0.001 | 0.000 | 0.032 | 0.000 |
| Genomator (H = 1) | 0.003 | 0.000 | 0.013 | 0.750 |
| Genomator (H = 10) | 0.026 | 0.020 | 0.018 | 0.789 |
| Genomator (H = 50) | 0.022 | 0.057 | 0.043 | 0.813 |

**Supplementary Table 2.** Kinship replay index (KRI) submetrics for the 10,000 SNPs panel dataset (chromosome 15).

| **Model** | **MIA_AUC_** | **r_mia_** | **U** | **U_0_** | **r_uniq_** |
| --- | --- | --- | --- | --- | --- |
| GAN | 0.483 | 0.000 | 0.664 | 0.539 | 0.271 |
| RBM | 0.500 | 0.001 | 0.905 | 0.539 | 0.794 |
| Genomator (H = 1) | 0.520 | 0.039 | 0.008 | 0.538 | 0.000 |
| Genomator (H = 10) | 0.512 | 0.024 | 0.005 | 0.538 | 0.000 |
| Genomator (H = 50) | 0.518 | 0.035 | 0.003 | 0.538 | 0.000 |

**Supplementary Table 3.** Trait linkage index (TLI) submetrics for the 10,000 SNPs panel dataset (chromosome 15).

| **Model** | **ρ_q_** | **r_ρ_** | **A_adv_** | **r_A_** |
| --- | --- | --- | --- | --- |
| GAN | 1.230 | 0.000 | 0.723 | 0.000 |
| RBM | 1.259 | 0.000 | 0.806 | 0.000 |
| Genomator (H = 1) | 0.569 | 0.431 | 0.323 | 0.353 |
| Genomator (H = 10) | 0.817 | 0.183 | 0.421 | 0.158 |
| Genomator (H = 50) | 0.949 | 0.052 | 0.523 | 0.000 |

**Supplementary Table 4.** Proximity leakage index (PLI) submetrics for the 65,535 SNPs panel dataset (chromosome 1).

| **Model** | **r_replay_** | **r_IKE_** | **r_HAP_** | **r_SPEC_** |
| --- | --- | --- | --- | --- |
| GAN | 0.000 | 0.000 | 0.000 | 0.541 |
| RBM | 0.002 | 0.000 | 0.000 | 0.000 |
| Genomator (H = 1) | 0.000 | 0.000 | 0.034 | 0.441 |
| Genomator (H = 10) | 0.000 | 0.023 | 0.055 | 0.488 |
| Genomator (H = 50) | 0.000 | 0.093 | 0.082 | 0.560 |

**Supplementary Table 5.** Kinship replay index (KRI) submetrics for the 65,535 SNPs panel dataset (chromosome 1).

| **Model** | **MIA_AUC_** | **r_mia_** | **U** | **U_0_** | **r_uniq_** |
| --- | --- | --- | --- | --- | --- |
| GAN | 0.502 | 0.003 | 0.636 | 0.719 | 0.000 |
| RBM | 0.514 | 0.028 | 0.875 | 0.719 | 0.555 |
| Genomator (H = 1) | 0.535 | 0.070 | 0.026 | 0.718 | 0.000 |
| Genomator (H = 10) | 0.526 | 0.052 | 0.019 | 0.718 | 0.000 |
| Genomator (H = 50) | 0.540 | 0.079 | 0.021 | 0.718 | 0.000 |

**Supplementary Table 6.** Trait linkage index (TLI) submetrics for the 65,535 SNPs panel dataset (chromosome 1).

| **Model** | **PLI** | **KRI** | **TLI** | **α/β** | **PRISM-G** |
| --- | --- | --- | --- | --- | --- |
| Safe binomial sampler | 0.005 | 0.108 | 0.012 | 0.008 | 0.000 |
| Leaky copycat | 1.000 | 0.206 | 1.000 | 0.748 | 100.000 |

**Supplementary Table 7.** PRISM-G scores and metrics for calibration anchors (10,000 SNPs panel dataset).

| **Model** | **PLI** | **KRI** | **TLI** | **α/β** | **PRISM-G** |
| --- | --- | --- | --- | --- | --- |
| Safe binomial sampler | 0.000 | 0.000 | 0.019 | 0.013 | 0.000 |
| Leaky copycat | 1.000 | 0.003 | 1.000 | 0.749 | 100.000 |

**Supplementary Table 8.** PRISM-G scores and metrics for calibration anchors (65,535 SNPs panel dataset).

| **Model** | **PLI** | **KRI** | **TLI** | **R_raw_** |
| --- | --- | --- | --- | --- |
| GAN | 0.301 | 0.435 | 0.271 | 0.272 |
| RBM | 0.345 | 0.018 | 0.794 | 0.568 |
| Genomator (H = 1) | 0.828 | 0.750 | 0.039 | 0.269 |
| Genomator (H = 10) | 0.808 | 0.789 | 0.024 | 0.261 |
| Genomator (H = 50) | 0.733 | 0.813 | 0.035 | 0.254 |

**Supplementary Table 9.** PRISM-G components for 10,000 SNPs panel dataset (chromosome 15).

| **Model** | **PLI** | **KRI** | **TLI** | **R_raw_** |
| --- | --- | --- | --- | --- |
| GAN | 0.000 | 0.541 | 0.003 | 0.051 |
| RBM | 0.000 | 0.000 | 0.555 | 0.412 |
| Genomator (H = 1) | 0.431 | 0.441 | 0.070 | 0.183 |
| Genomator (H = 10) | 0.183 | 0.488 | 0.051 | 0.118 |
| Genomator (H = 50) | 0.052 | 0.560 | 0.079 | 0.111 |

**Supplementary Table 10.** PRISM-G components for 65,535 SNPs panel dataset (chromosome 1).

| **Model** | **PRISM-G** | **SD** | **LCI (95%)** | **UCI (95%)** |
| --- | --- | --- | --- | --- |
| RBM | 70.368 | 8.920 | 47.738 | 77.311 |
| Genomator (H = 1) | 45.531 | 15.041 | 31.758 | 80.729 |
| Genomator (H = 10) | 44.104 | 14.842 | 30.976 | 79.199 |
| Genomator (H = 50) | 41.576 | 13.102 | 31.074 | 73.304 |
| GAN | 35.312 | 1.070 | 33.589 | 37.970 |

**Supplementary Table 11.** PRISM-G scores for 10,000 SNPs panel dataset (chromosome 15; default hyperparameters; 1,000 bootstraps). SD: standard deviation; LCI: lower confidence interval; UCI; upper confidence interval.

| **Model** | **PRISM-G** | **SD** | **LCI (95%)** | **UCI (95%)** |
| --- | --- | --- | --- | --- |
| RBM | 42.763 | 7.404 | 22.677 | 50.267 |
| Genomator (H = 1) | 26.534 | 4.638 | 22.144 | 39.300 |
| Genomator (H = 10) | 13.756 | 1.669 | 11.001 | 18.542 |
| Genomator (H = 50) | 10.111 | 2.615 | 6.065 | 14.812 |
| GAN | 2.102 | 2.223 | 0 | 6.682 |

**Supplementary Table 12.** PRISM-G scores for 65,535 SNPs panel dataset (chromosome 1; default hyperparameters; 1,000 bootstraps). SD: standard deviation; LCI: lower confidence interval; UCI; upper confidence interval.

| **Model** | **PRISM-G** | **SD** | **LCI (95%)** | **UCI (95%)** |
| --- | --- | --- | --- | --- |
| RBM | 69.761 | 5.832 | 56.027 | 74.412 |
| Genomator (H = 1) | 48.910 | 10.372 | 39.747 | 71.715 |
| Genomator (H = 10) | 47.313 | 10.259 | 38.332 | 69.986 |
| Genomator (H = 50) | 44.165 | 9.080 | 36.369 | 64.486 |
| GAN | 35.392 | 0.716 | 34.091 | 36.944 |

**Supplementary Table 13.** PRISM-G scores 10,000 SNPs panel dataset (chromosome 15; grid search results). SD: standard deviation; LCI: lower confidence interval; UCI; upper confidence interval.

| **Model** | **PRISM-G** | **SD** | **LCI (95%)** | **UCI (95%)** |
| --- | --- | --- | --- | --- |
| RBM | 42.034 | 3.837 | 30.477 | 45.554 |
| Genomator (H = 1) | 27.535 | 2.840 | 24.594 | 36.179 |
| Genomator (H = 10) | 13.847 | 1.153 | 12.008 | 17.148 |
| Genomator (H = 50) | 9.628 | 1.089 | 7.159 | 11.239 |
| GAN | 1.204 | 0.880 | 0 | 2.740 |

**Supplementary Table 14.** PRISM-G scores and metrics for 65,535 SNPs panel dataset (chromosome 1; grid search results). SD: standard deviation; LCI: lower confidence interval; UCI; upper confidence interval.

| **Model** | **Mean diff.** | **LCI (95%)** | **UCI (95%)** | **P-value** |
| --- | --- | --- | --- | --- |
| GAN - RBM | -34.370 | -39.491 | -13.395 | 0 |
| GAN - Genomator (H = 1) | -13.519 | -35.084 | -4.658 | 0 |
| GAN - Genomator (H = 10) | -11.922 | -33.348 | -3.309 | 0 |
| Genomator (H = 1) - Genomator (H = 10) | 1.597 | 1.357 | 1.855 | 0 |
| Genomator (H = 10) - Genomator (H = 50) | 3.149 | 1.863 | 5.554 | 0 |
| Genomator (H = 1) - Genomator (H = 50) | 4.746 | 2.329 | 7.319 | 0 |
| GAN - Genomator (H = 50) | -8.773 | -27.825 | -1.458 | 0 |
| RBM - Genomator (H = 50) | 25.597 | -8.413 | 37.644 | 2.82E-01 |
| RBM - Genomator (H = 10) | 22.448 | -13.933 | 35.616 | 3.88E-01 |
| RBM - Genomator (H = 1) | 20.851 | -15.668 | 34.185 | 3.96E-01 |

**Supplementary Table 15.** Permutation test 10,000 SNPs panel dataset (chromosome 15; grid search results). LCI: lower confidence interval; UCI; upper confidence interval.

| **Model** | **Mean diff.** | **LCI (95%)** | **UCI (95%)** | **P-value** |
| --- | --- | --- | --- | --- |
| GAN - RBM | -40.830 | -43.958 | -29.980 | 0 |
| GAN - Genomator (H = 1) | -26.331 | -34.869 | -22.702 | 0 |
| GAN - Genomator (H = 10) | -12.643 | -15.969 | -11.287 | 0 |
| GAN - Genomator (H = 50) | -8.424 | -8.717 | -7.159 | 0 |
| RBM - Genomator (H = 10) | 28.186 | 13.949 | 32.206 | 0 |
| RBM - Genomator (H = 50) | 32.406 | 21.874 | 35.397 | 0 |
| Genomator (H = 1) - Genomator (H = 10) | 13.687 | 11.416 | 19.031 | 0 |
| Genomator (H = 1) - Genomator (H = 50) | 17.907 | 14.069 | 26.905 | 0 |
| Genomator (H = 10) - Genomator (H = 50) | 4.219 | 2.654 | 7.921 | 0 |
| RBM - Genomator (H = 1) | 14.499 | -5.065 | 20.340 | 1.46E-01 |

**Supplementary Table 16.** Permutation test for 65,535 SNPs panel dataset (chromosome 1; grid search results). LCI: lower confidence interval; UCI; upper confidence interval.

**Supplementary Figures**

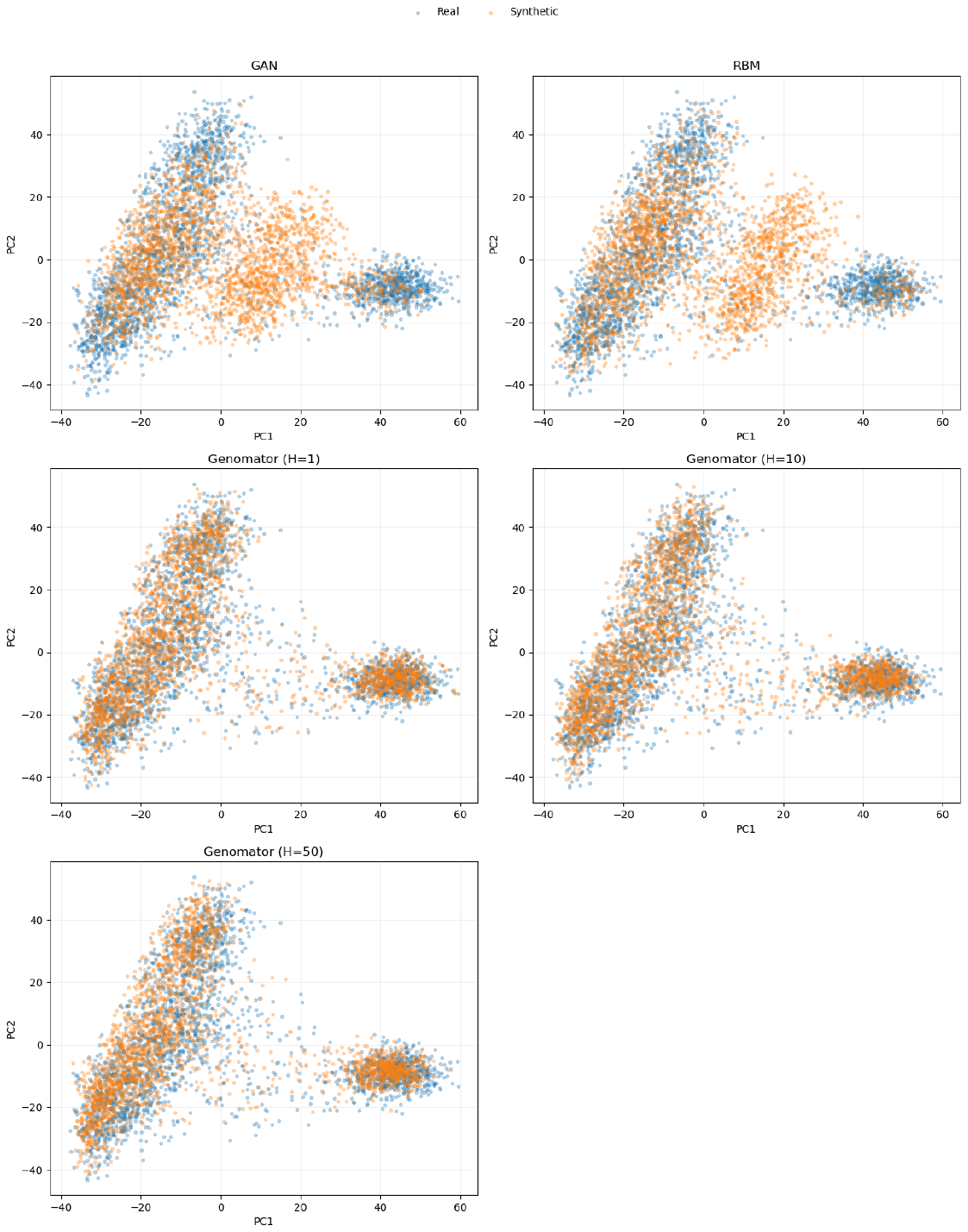

**Figure 1.** PCA of real and synthetic genomes (Chromosome 15; 10,000 SNPs). PCA was fitted on the real dataset and synthetic samples were projected onto the same PCA space. Blue points represent real samples and orange points represent synthetic samples generated by different models (GAN, RBM, and Genomator with H = 1, 10, 50). The overlap indicates how well each model reproduces the population structure.

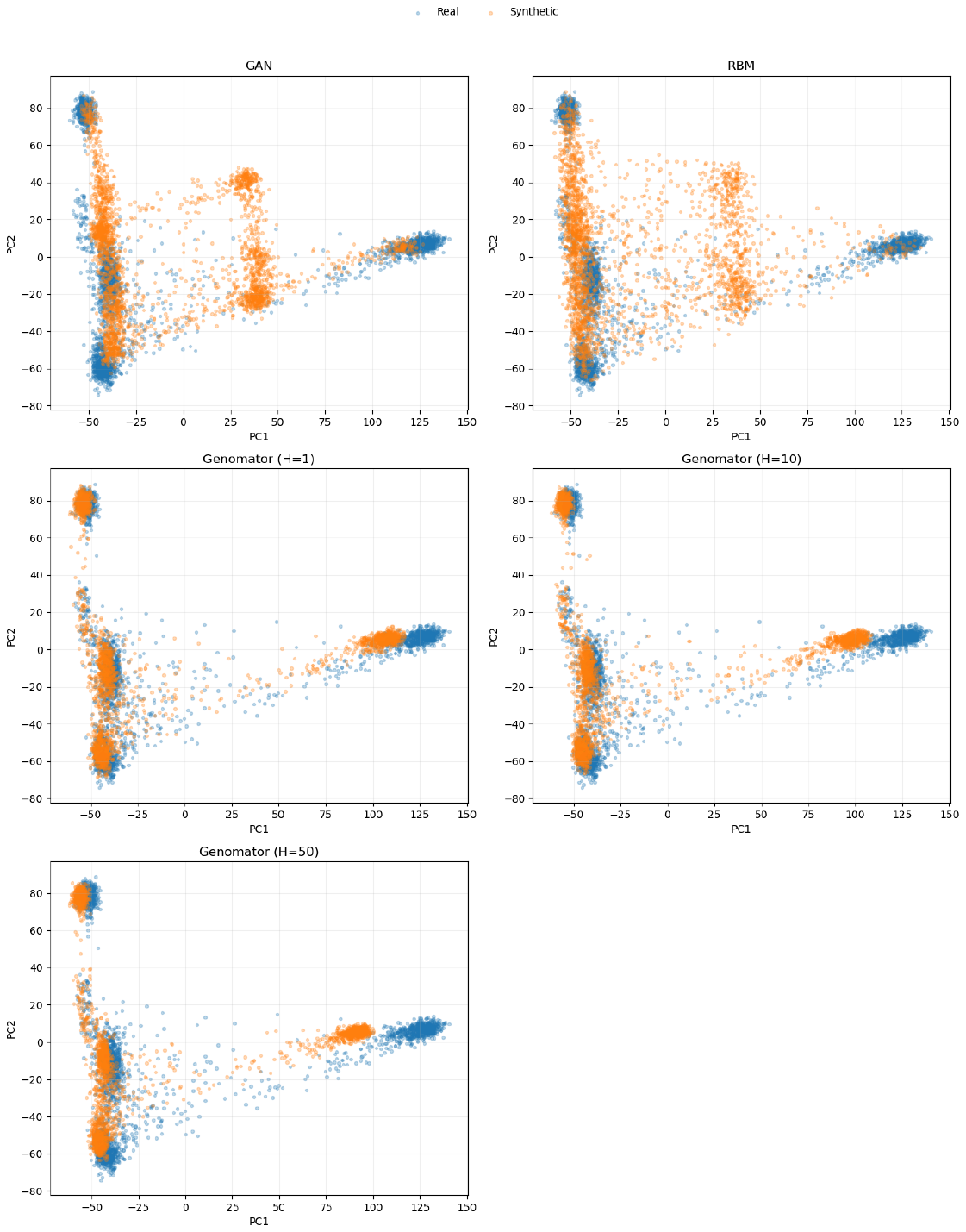

**Figure 2.** PCA of real and synthetic genomes (Chromosome 1; 65,535 SNPs). PCA was fitted on the real dataset and synthetic samples were projected onto the same PCA space. Blue points represent real samples and orange points represent synthetic samples generated by different models (GAN, RBM, and Genomator with H = 1, 10, 50). The overlap indicates how well each model reproduces the population structure.
